# Midbrain dopamine neuronal activity modulates splenic immunity through a brain-to-body circuit

**DOI:** 10.1101/2024.12.05.625471

**Authors:** Adithya Gopinath, Alfonso junior Apicella, Alice Bertero, Leah T. Phan, Emily Miller, Jordan Seibold, Shannon C. Wall, Jordan Follett, Aidan Smith, Elijah Freedman, Lauryn Hornstein, Logan Wood, Hannah Ciupe, Diya Behal, Arianna Grillo, Delaney Booth, Sabrina Campos, Nirvana Soundara, Daniel L. Purich, Pedro Trevizan-Baú, Leah R. Reznikov, Nikhil Urs, Olga Guryanova, Mathew J. Farrer, Daniel W. Wesson, Habibeh Khoshbouei

## Abstract

Midbrain dopamine neurons are well-known to shape central nervous system function, yet there is growing evidence for their influence on the peripheral immune system. Here we demonstrate that midbrain dopamine neurons form a circuit to the spleen via a multi-synaptic pathway from the dorsal vagal complex (DVC) through the celiac ganglion. Midbrain dopamine neurons modulate the activity of D1- and D2-like dopamine receptor-expressing DVC neurons. *In vivo* activation of midbrain dopamine neurons induced dopamine release in the DVC and increased immediate early gene expression in both the DVC and celiac ganglion. Activation of this midbrain-to-spleen circuit reduced spleen weight and decreased naïve CD4^+^ T-cell populations without affecting total T-cell numbers. These findings unveil a functional midbrain- DVC-celiac ganglion-spleen pathway, through which midbrain dopamine neurons modulate splenic immunity. These novel insights into the neural regulation of the immune system have important implications for diseases involving altered dopamine neurotransmission and highlight potential targets for immunotherapeutic interventions.

## Introduction

While substantial progress has been made in unraveling body-to-brain communication^1^, the reverse pathway, how the brain influences peripheral organs, particularly through dopamine signaling, remains unclear. Emerging evidence highlights that midbrain dopamine neurons exert significant influence on peripheral systems, particularly in the context of neuroimmune interactions, but the circuits connecting them to peripheral immune organs remain largely undefined. Zhu and colleagues^2^, for instance, identified a functional link between central nervous system (CNS) pain processing and splenic immunity, suggesting that neural pathways involved in immune regulation extend well beyond the brain’s immediate environment to influence key organs, such as the spleen. Such findings imply that midbrain dopaminergic neurons may play a previously uncharacterized role in orchestrating peripheral immune responses. Indeed, degeneration of midbrain dopamine neurons is closely associated with changes in peripheral immune dysfunction believed to contribute to disease progression^3-5^. Similarly, psychostimulant-induced activation of midbrain dopamine neurons regulates peripheral immunity and influence wound healing^6-11^. Moreover, the well-characterized connection between peripheral inflammation and infection, manifested symptomatically as anhedonia and reduced motivation^12-14^, strongly suggest a link between dopamine neurotransmission and peripheral immunity. Despite such findings, the specific circuits through which midbrain dopamine neurons communicate with peripheral immune organs remain unclear.

By connecting the brainstem to multiple peripheral organs including heart, lung, gut, and spleen^15-18^, the vagus nerve is one of the most characterized routes for brain-body communication. While body-to-brain communication through vagal efferents has been extensively studied, the corresponding brain-to-periphery circuits that use dopamine signaling to mediate the brain’s control over immune functions remain to be defined. The dorsal vagal complex (DVC) plays a pivotal role in modulating immune function through the cholinergic anti-inflammatory pathway^19-21^. The DVC includes the dorsal motor nucleus of the vagus (DMV), the area postrema (AP), and the nucleus of the solitary tract (NTS) and is optimally positioned to transmit signals from midbrain dopamine neurons to peripheral organs. Prior work revealed that midbrain neurons innervate the DVC and separately that midbrain neuron output appears to influence colon motility^22,23^. In addition, vagal innervation of peripheral organs including the spleen contribute to both inflammation and its’ resolution^24-28^. While these advances have strongly supported the existence of a brain-to-periphery system which can influence both peripheral organs and immune system function, the specific role of dopamine in the modulation of splenic immunity is unclear.

The spleen is innervated by both cholinergic^29-31^ and catecholaminergic^32-35^ fibers and is essential for survival, with roles in the production of blood cells and immune function. Both inputs modulate splenic function, immune cell populations and the broader peripheral immune landscape. Here we used a combination of viral approaches to map a midbrain dopamine neurons→DVC→ spleen circuit, focusing on how midbrain dopamine neuron activity modulates splenic immune phenotype. We demonstrate that midbrain dopamine neurons project to the DVC, can activate dopaminergic receptors expressed on DVC neurons, and that their activity is transmitted through the celiac ganglia to reach the spleen. Furthermore, we show that the activation of this circuit alters peripheral immune function, including spleen weight and splenic T-cell populations. Notably, transection of the cervical vagus nerve abolishes the impact of midbrain dopamine neuron activation on splenic immunity, confirming the essential role of vagal signaling in this brain-to-spleen pathway. Collectively, our findings provide novel insights into the mechanisms underlying the regulation of peripheral immunity by midbrain dopamine neurons, integrating prior observations into a cohesive circuit model, casting new light on the interplay between central dopamine signaling and peripheral immunity, and taking us closer to the root causes of disorders characterized by dysregulated CNS dopamine signaling.

## Results and Discussion

### Anterograde mapping of midbrain dopaminergic neurons confirms dopaminergic projections to the dorsal vagal complex

Prior work established CNS circuity linking midbrain with the DVC^1,2,22,23,36-38^. Here we built upon these reports and performed two initial viral-tracing based studies to identify 1) are midbrain neuron inputs to the DVC dopaminergic and 2) are they monosynaptic?

To address the first question we utilized DAT-Cre mice, which express Cre recombinase under the control of the dopamine transporter (DAT)^39^, and injected AAV2-Ef1a-DIO-EYFP into the midbrain (Figure 1A). This allowed for EYFP expression in tyrosine hydroxylase+ (TH^+^) dopamine neuron cell bodies in the midbrain as well as their axons in the median forebrain bundle (Figures 1B & 1C and Supplementary Figure 1). As the gold standard for cholinergic neurons, choline acetyltransferase (ChAT) co-staining was used to identify the DVC (Figure 1D). As expected, EYFP^+^ fibers were detected throughout the DVC (Figures 1D-E), including in the DMV and NTS, showing midbrain dopaminergic neurons send projections to the DVC.

**Figure 1.**
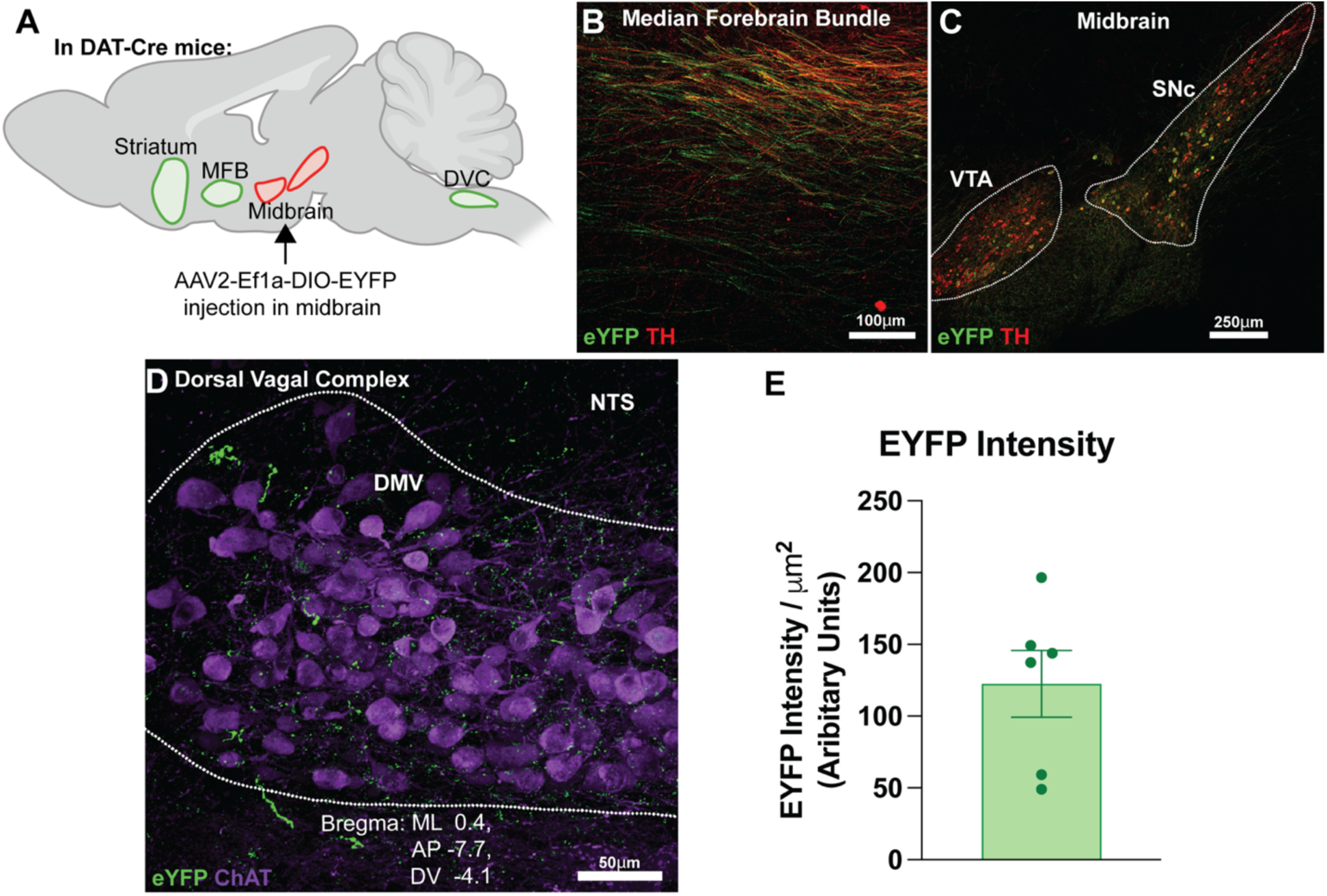
Anterograde mapping of midbrain dopaminergic neurons confirms dopaminergic projections to the dorsal vagal complex. **Panel A:** Anatomic location of injection of AAV2-Ef1a-DIO-EYFP into the midbrain of DAT^IRES^-Cre mice, followed by three-week expression period. **Panel B:** EYFP expression in positive control region, the median forebrain bundle (MFB). **Panel C:** EYFP expression at the injection site in the midbrain. Note also that tyrosine hydroxylase (TH) co-staining confirmed the fidelity of expression in dopaminergic projections from the midbrain in B and C. **Panel D:** EYFP+ projections surrounding ChAT+ DMV neurons in the DVC, suggesting monosynaptic projections from the midbrain to DVC. **Panel E:** EYFP intensity per square millimeter (expressed as arbitrary fluorescence units). *Data representative of n=6 independent experiments*.

We next investigated if these midbrain dopaminergic neurons form synapses within the DVC. Using a complementary viral tracing approach (Figure 2A), we injected DAT-Cre mice with AAV1-hSyn-Flex-mGFP-2A-Synaptophysin-mRuby into the midbrain. This drives expression of the Synaptophysin-mRuby fusion protein in dopaminergic synapses that originate form midbrain neurons^40^. We next inspected the DVC for mRuby^+^ puncta. mRuby^+^ puncta were confirmed at the injection site in the midbrain as well as in known projection areas, such as the striatum (Figure 2B & 2C, Supplemental Figure 2). Importantly, mRuby^+^ puncta were observed throughout the DVC, including in the DMV and NTS (Figure 2D-E), indicating that DAT^+^ midbrain dopamine neurons form synapses within the DVC and its subnuclei.

**Figure 2.**
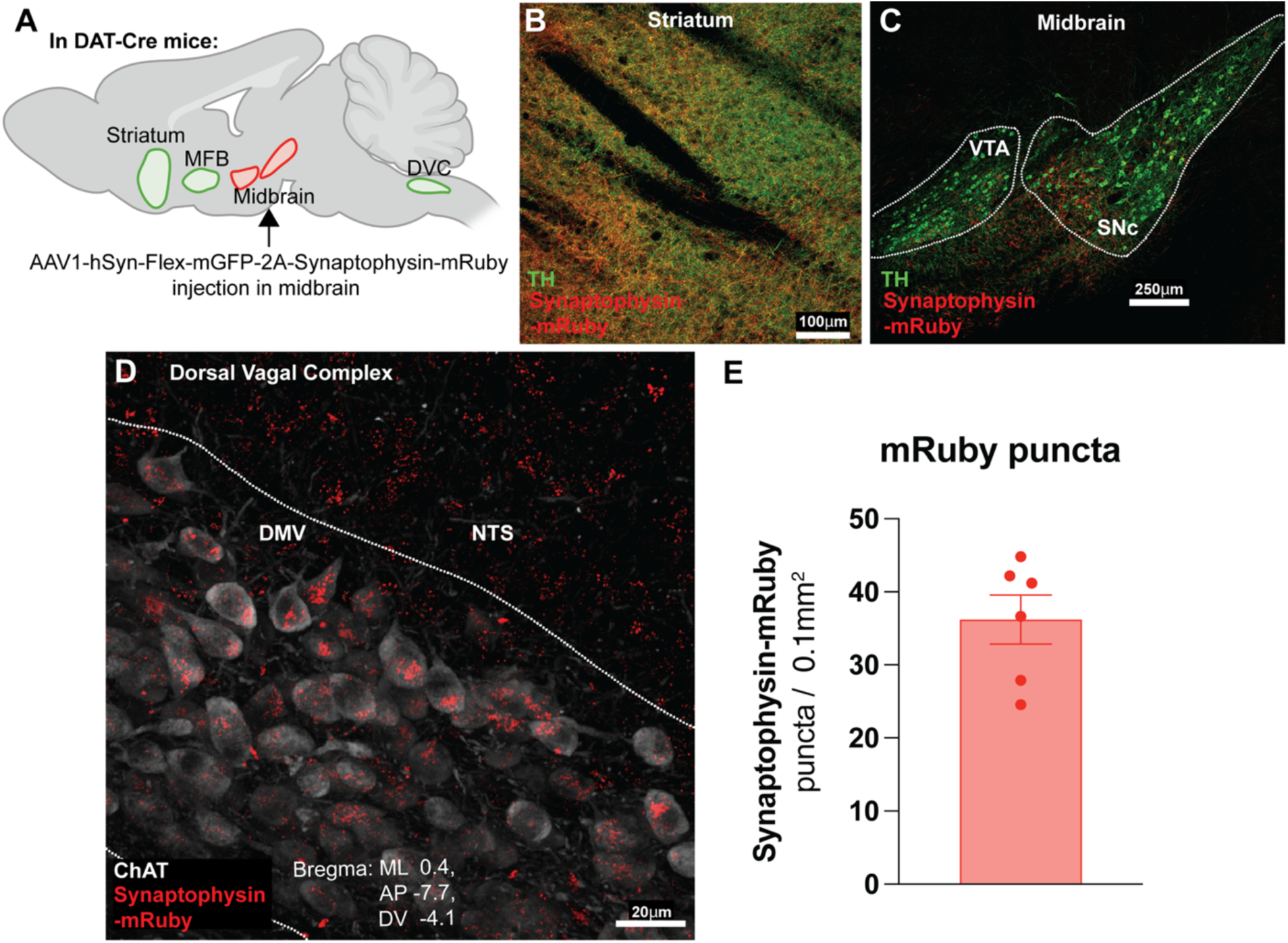
DAT+ midbrain neurons synapse throughout the dorsal vagal complex. **Panel A:** Anatomic locations in brains of DAT-Cre mice receiving midbrain injections of AAV1-hSyn-Flex-mGFP-2A-Synaptophysin-mRuby. Three weeks after midbrain injection, the resulting fluorescence patterns indicate that midbrain dopamine neurons not only send projections to the DVC but also make synapses in the DVC. **Panel B:** synaptophysin-mRuby+ puncta in control regions of striatum. **Panel C:** synaptophysin-mRuby+ puncta at the injection site in the midbrain. [Panels B and C also show tyrosine hydroxylase (TH) co-staining to confirm expression in dopaminergic projections from the midbrain.] **Panel D:** Presence of synaptophysin-mRuby+ puncta throughout DVC, identified by presence of choline acetyltransferase+ (ChAT+) neurons. **Panel E:** Quantification of synaptophysin-mRuby+ puncta. *Data representative of n=6 independent experiments*.

### DVC neurons express D1- and D2-like receptors

The above findings support a two-part hypothesis: 1) changes in midbrain dopamine neuron activity and dopamine release modulate DVC neuronal activity; and 2) this modulation, in turn, regulates peripheral systems downstream of the DVC, and may comprise a previously unrecognized component of the midbrain dopamine neuron-to-peripheral immunity circuit. As noted above, choline acetyltransferase (ChAT) is a reliable marker to help identify the DVC. To establish a receptor-based foundation for testing our hypotheses, we performed whole-cell patch-clamp recordings in ChAT-GFP mice, targeting GFP-expressing ChAT^+^ neurons, the major parasympathetic output population^20,41^, as well as GFP-negative neurons in the DVC. This experimental approach allows for an assessment the effects of dopamine on the intrinsic excitability and firing properties of the neurons identified. Whole-cell patch-clamp recordings were performed on coronal brainstem slices in the presence of synaptic blockers for GABAergic and glutamatergic transmission, enabling isolation of the direct, postsynaptic actions of dopamine (Figure 3A). Firstly, since GFP expression might highlight only a subset of ChAT neurons in the DVC and cause misleading identification of DVC neurons’ identity, we compared the intrinsic properties of GFP-positive (N=22) and GFP-negative (N=19) neurons, to determine if they belong to two separate classes of neurons. The vast majority of GFP-positive neurons (77%, 17/22) showed spontaneous firing activity when recorded in cell-attached or current-clamp configuration (Figure 3B), a common feature of ChAT-expressing neurons in other brain regions^42^ and of DVC neurons^43^. GFP-negative neurons, on the other hand, do not display such behavior. The two populations have comparable passive properties, such as input resistance (Ri GFP^+^ 639.6 ± 100.5 MΩ, Ri GFP^-^ 878.9 ± 80.3 MΩ, p=0.076) and membrane time constant (tau, GFP^+^ 0.46 ± 0.03 msec, GFP^-^ 0.36 ± 0.03 msec) (Figure 3C), and their voltage threshold to fire an action potential (V threshold, GFP^+^ -32.1 ±1.5 mV, GFP^-^ -31.9 ±1.6 mV, p=0.92, Figure 3C). However, they display a wide array of differences in their active properties, such as current threshold (I threshold, GFP^+^ 81.8 ± 8.4 pA, GFP^-^ 50 ± 0 pA, p=0.001), action potential height (V height, GFP^+^ 90.6 ± 2.3 mV, GFP^-^ 59.0 ± 2.2 mV, p=0.0008), action potential halfwidth (AP halfwidth, GFP^+^ 0.73 ± 0.04 msec, GFP^-^ 0.55 ± 0.2 msec, p=0.0003), the relationship between injected current and AP frequency (f/I slope, GFP^+^ 0.08 ± 0.01 Hz/pA, GFP^-^ 0.25 ± 0.05 Hz/pA, p=0.001) and spike frequency adaptation ratio (SFA ratio GFP^+^ 1.07 ± 0.011, GFP^-^ 0.59 ±0.05, p=0.0006) (Figure 3D). This confirms that GFP-positive and GFP-negative neurons in the DVC are two separate neuronal populations with essentially different firing properties.

**Figure 3.**
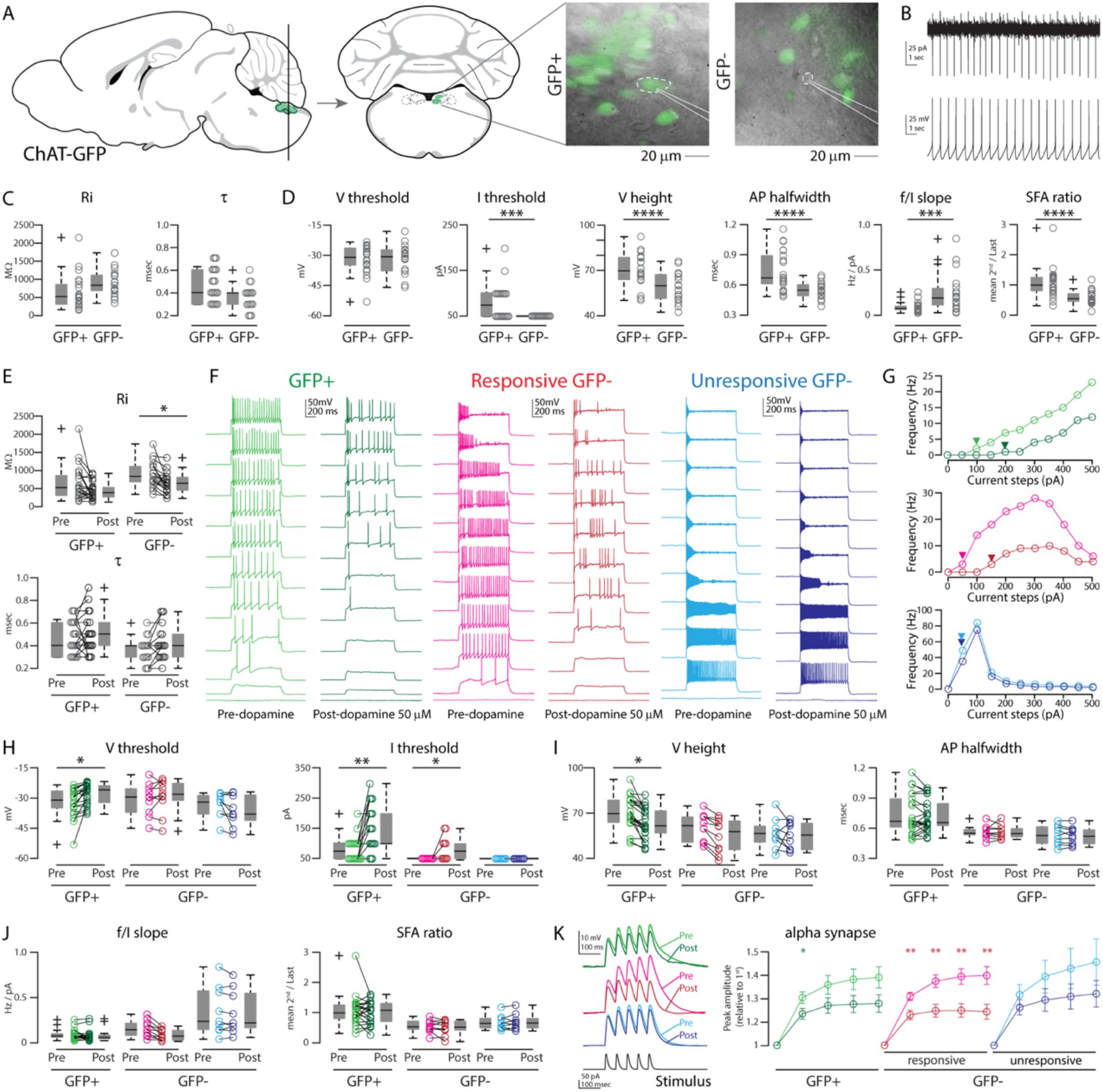
DVC neurons express D1-like and D2-like receptors. **Panel A:** DVC neurons of adult Chat-GFP mice were recorded, using whole-cell patch clamp. Left: Anatomic drawing of brain (sagittal), showing region of interest (green shading). Vertical line represents the coronal plane(s) collected for patching, as shown in middle panel. Right: Position of the patching pipette on a GFP-positive and a GFP-negative neuron in the DVC. **Panel B**: GFP-expressing neurons are tonically active: they spontaneously fire action potentials in resting conditions, both in cell-attach (top trace) and in whole-cell current-clamp configuration (bottom trace). **Panel C**: GFP-expressing and GFP-negative neurons display similarities in intrinsic properties such as input resistance (Ri), membrane time constant (τ). **Panel D**: active properties of GFP-expressing and GFP-negative neurons in the DVC: V_threshold_, the membrane voltage threshold to generate an action potential; *I*_threshold_, the minimal current step to elicit an action potential (rheobase); *V*_Height_ the total amplitude of the action potential (AP); action potential halfwidth (AP_halfwidth_ the width of the first action potential at threshold, measured at half the peak amplitude); f/I slope, the relationship between AP frequency and increasing depolarizing currents applied; SFA, spike frequency adaptation, measured as the ratio between the second and last instantaneous frequency of a train of action potentials upon depolarizing currents. *** = p<0.005, **** = p<0.001 (two tailed t test). **Panel E:** Summary boxplots of passive properties of GFP-positive (GFP+, shades of green, N=22) and GFP-negative (GFP-, shades of red, N=19) neurons in the DVC, before (“Pre”, lighter circles) and after (“Post”, darker circles) bath administration of 50 μM dopamine. **Panel F:** Representative traces from the three populations identified, showing the membrane responses to increasing depolarizing current steps (from 0 pA to 500 pA in 50 pA steps). The lighter traces were recorded in baseline conditions, holding the neurons at -70mV in absence of dopamine, while the darker traces were recorded 4 minutes after bath-application of 50 μM dopamine. **Panel G**: Quantification of the firing rate (frequency of action potentials, Hz) in response to depolarizing currents (pA) of each cell type in Panel-E, same color coding: GFP+ neuron before (light green) and after dopamine application (dark green); responsive GFP-neuron before (magenta) and after dopamine application (red); unresponsive GFP-neuron before (light blue) and after dopamine application (dark blue). Arrowheads indicate rheobases. **Panel H**: Quantification of V threshold and I threshold for the three populations identified, pre (light colored circles) and post bath application of dopamine (dark colored circles). **Panel I**: Quantification of V height and AP halfwidth for the three populations identified, pre (light colored circles) and post bath application of dopamine (dark colored circles). **Panel J:** Quantification of f/I slope and SFA ratio for the three populations identified, pre (light colored circles) and post bath application of dopamine (dark colored circles). **Panel K**: Left: three representative alpha-synapse responses from the populations analyzed pre (lighter traces) and post (darker traces) bath application of dopamine. Right: population analysis of synaptic integration, expressed as a fold increase on the first of five EPSP-like inputs before (lighter traces) and after (darker traces) bath application of dopamine, showing a significant effect on GFP+ and responsive GFP-neurons in the DVC. * = p <0.05; ** = p<0.01 (two tailed t-test comparing before and after bath application of dopamine).

We next examined how dopamine modulates the intrinsic properties of GFP-positive (N=22) and GFP-negative (N=19) DVC neurons. After determining their baseline intrinsic properties, we applied 50 μM dopamine to the ACSF bath and reassessed the neurons’ properties, for comparison. Overall, dopamine did not change the passive membrane properties of GFP-positive neurons (Figure 3E) but affected the input resistance of GFP-negative neurons, which showed a significant decrease (Pre dopamine: 878.89 ± 80.26 MΩ; Post dopamine: 656.37 ± 63.43 MΩ; p=0.036, Figure 3E). Based on the effect of the firing properties (Figure 3F, Supplementary Table 1), we could identify three main neuronal groups: GFP-expressing ChAT neurons (N=22) and GFP-negative neurons (N=10/19) with an overall decrease in firing activity and increase in rheobase, and a subset of GFP-negative neurons (N=9/19) unresponsive to dopamine, as shown in three representative examples in Figure 3F, whose f/I curve, representing their changes in firing pattern, demonstrate the criteria for classification (Figure 3G, Supplementary Table 1). The two responsive DVC populations show significant changes in membrane and current thresholds (Figure 3H, Supplementary Table 1) and V height, but not in AP halfwidth, f/I slope and SFA ratio (Figure 3I-J, Supplementary Table 1). This behavior is compatible with D2 receptor expression ^44,45^. It is worth noting that a very small fraction of GFP-expressing ChAT-positive neurons showed an increased F/I slope (Figure 3I, F/I slope, N=2/22, Supplementary Table 1) without changes in rheobase. Such behavior suggests enhanced excitability following dopamine administration, consistent with the expression of D1 receptors^23,46^. We also tested if dopamine influences the synaptic integration of DVC neurons by injecting a train of 5 EPSP-like waveform at 20Hz (α-EPSPs) into the soma via the patch pipette. We found that, after bath application of dopamine, the three populations showed different alterations of synaptic integration, with responsive GFP-negative neurons showing the most significant reduction in temporal summation (Figure 3K, Supplementary Table 1). Overall, these findings indicate that both ChAT-positive and ChAT-negative neurons in the DVC respond to dopamine, exhibiting changes in intrinsic excitability, firing patterns, and synaptic integration properties.

### Stimulation of midbrain dopamine neurons induces dopamine release in the DVC

We next tested the hypothesis that stimulation of midbrain dopamine neurons causes dopamine release in the DVC. We injected AAV2-hSyn-DIO-hM3D(Gq)-mCherry (encoding excitatory Gq-coupled DREADDs) into the midbrain of DAT-Cre mice. Three weeks post-injection, the Cre-mediated expression of Gq-DREADD-mCherry was verified in the somatic (midbrain) and terminal regions (striatum and DVC) (Figure 4C, Supplementary Figure 3).

**Figure 4.**
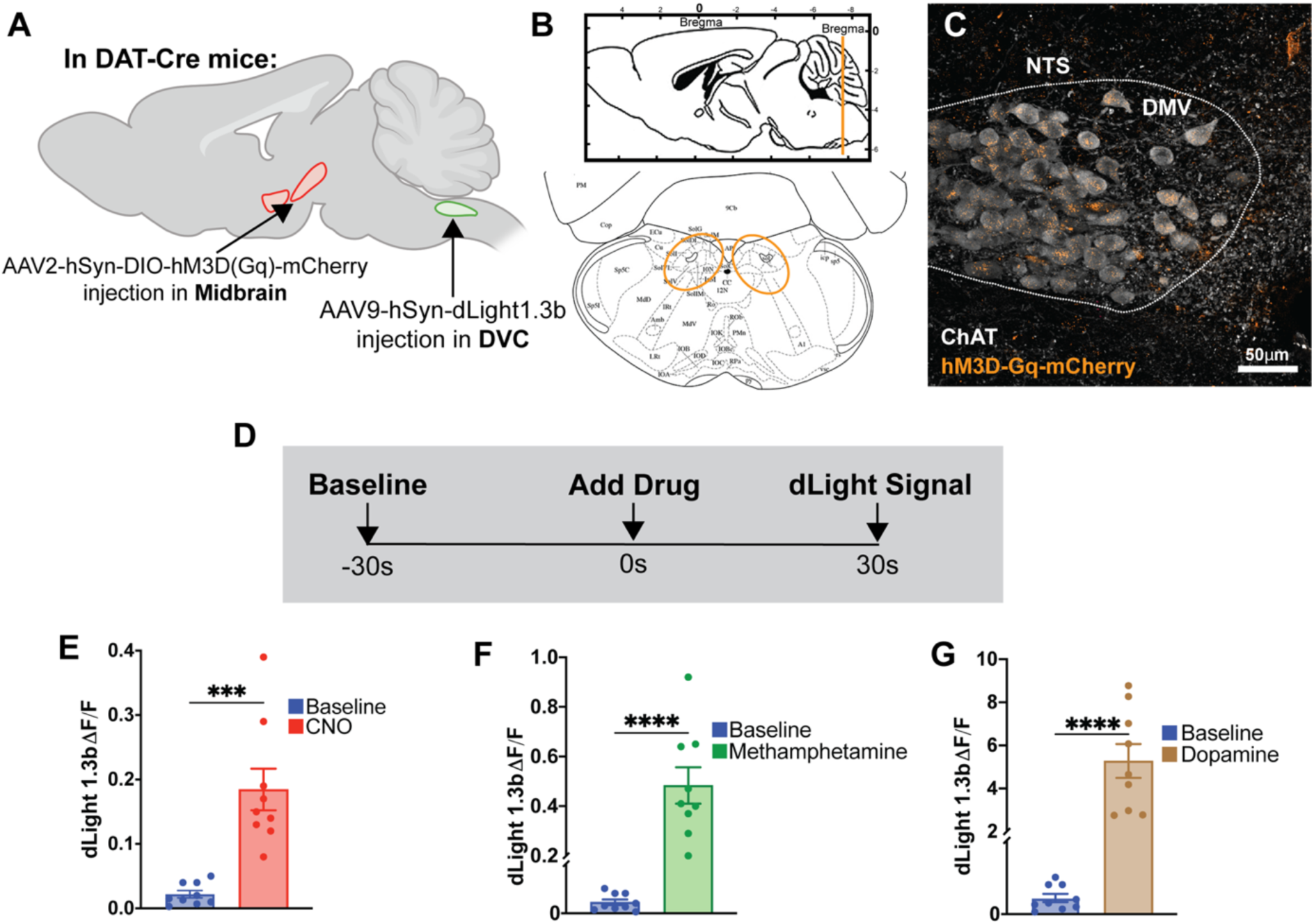
Midbrain dopamine neuron projections expressing excitatory Gq-DREADD receptors in the DVC release dopamine upon stimulation with CNO, and upon stimulation with methamphetamine, indicating functional projections from midbrain DAT+ neurons to DVC. **Panel A:** Anatomic midbrain locations of DAT-Cre mice receiving AAV2-DIO-hM3D(Gq)-DREADD-mCherry. In the same surgery, mice received AAV9-hSyn-dLight1.3b in the DVC, followed by expression for 3 weeks. **Panel B:** Coronal sections containing DVC (highlighted in orange) were obtained. **Panel C:** Co-staining for ChAT to identify DVC where DREADD expression in dopamine neuron terminals was verified (data representative of *n* = 3 biological replicates). **Panel D:** Live 250um coronal sections containing the DVC were collected and placed in a recovery chamber at 34 degrees C with carbogenation. After allowing for 30 minutes for rest and recovery, sections were transferred to an imaging chamber perfused with carbogenated ACSF. After collecting a fluorescence baseline (-30 sec) (Panel D-experimental design), drug perfusion was initiated (time zero). Peak signal (30 seconds) was assessed by subtracting baseline fluorescence from stimulated dLight1.3b fluorescence at the 30 second mark. **Panel E:** Stimulated release of dopamine from dopamine terminals originating in the midbrain upon bath perfusion of CNO (10 μM), resulting in a significant increase in dLight1.3b signal. **Panel F:** Additional incremental increase in DA release within the DVC, upon bath perfusion of methamphetamine (10 μM), as determined in separate experiments. **Panel G:** Positive control obtained with 100 μM dopamine (*n=4 independent biological replicates, with 2 technical replicates per experiment; ***p<0.005, ****p<0.0005, two-tailed unpaired Student’s T-test*).

In a separate cohort, we injected the same Gq-DREADD-encoding AAV into the midbrain and AAV9-hSyn-dLight1.3b into the DVC, leading to the expression of the high-affinity dopamine sensor dLight1.3b^47^ in the DVC (Figure 4A-B). Three weeks later, 250 μm coronal slices of the DVC were imaged with epifluorescence microscopy. For each experimental condition, pre-drug fluorescence signals (-30 seconds → 0 seconds) were subtracted from the signal peak after drug application (30 seconds) (Figure 4D) (Materials and Methods). We observed significantly increased dLight1.3b fluorescence after clozapine-N-oxide (CNO) application (Figure 4E), indicating that dopamine is released from Gq-DREADD-expressing midbrain-originating dopamine terminals in the DVC. DVC sections were also imaged at baseline and after perfusion with either 10 μM methamphetamine (Figure 4F), a well-characterized stimulator of dopamine release^48-50^, or 0.1 μM dopamine (Figure 4G) - another positive control. Both treatments significantly increased dLight1.3b fluorescence, confirming the specificity of dopamine release from dopaminergic terminals in the DVC. These findings support the conclusion that the activation of midbrain dopamine neurons induces dopamine release in the DVC, and suggests that dopaminergic projections from the midbrain regulate DVC activity.

### Anterograde multi-transsynaptic tracing reveals midbrain-to-spleen connectivity via celiac ganglion

The celiac ganglion, which receives projections from the DVC, acts as a relay to peripheral organs, including the liver, bone marrow and spleen^51-55^. Vagal afferents via the celiac ganglion regulate splenic immune functions^33,56-59^. Decades of research support the existence of the celiac ganglia-DVC circuit^60,61^, and with the midbrain dopamine neuron-to-DVC projections identified in this study, we propose a multi-synaptic link between midbrain dopamine neurons and the spleen. We used the Ai9 TdTomato Cre-reporter line^62^ and performed midbrain injections of pENN-AAV1-hSyn-Cre-WPRE-hGH to transduce Cre in an anterograde multi-transsynaptic manner^63^ (Figure 5A). Three weeks later, TdTomato^+^ cell bodies were observed in the DVC (Fig 5C). We also observed TdTomato expression in both the celiac ganglion and spleen (Figure 5D-E, Supplementary Figure 5). TdTomato^+^ expression in the spleen surrounded T-cell zones, identified by CD3^+^ T-cells (Figure 5E, Supplementary Figure 5). As that the spleen contains few neuronal cell bodies^64^, the TdTomato detected in the spleen likely reflects neuronal fibers innervating the spleen from the celiac ganglion, reinforcing the multi-synaptic midbrain-DVC-celiac ganglion-spleen projection pathway. Although these results suggest a midbrain-DVC-celiac ganglion-spleen pathway, they do not exclude the possibility that other direct or indirect connections exist which may also influence peripheral immunity^65,66^. DVC to the celiac ganglion connections likely reflect a composite of efferent (CNS-to-periphery) and afferent (periphery-to-CNS) activity, and would involve sympathetic spinal inputs to the celiac ganglion. In this study, our experiments are designed to interrogate the efferent arm of this pathway, namely the DVC → celiac ganglia → spleen axis. In addition, the efferent modulation within a midbrain–DVC–celiac ganglion–spleen circuit is likely to be accompanied by feedback through afferent pathways back to the CNS. Nevertheless, the midbrain-DVC-celiac ganglion-spleen pathway offers one mechanism by which midbrain dopamine neuron activity may influence splenic immune function. To investigate this possibility, we developed a stepwise experimental approach to assess 1) whether there is increased DVC neuronal activity when midbrain dopamine neurons are activated; 2) whether celiac ganglion neurons exhibit increased activity under the same conditions; 3) whether splenic immune cell populations are altered during dopamine neuron activation, and 4) whether perturbation of the midbrain-DVC-celiac ganglion-spleen pathway via vagus nerve transection abrogates these effects. Following this stepwise plan, we sought to investigate the functional impact of midbrain dopamine neuron activation at each junction of this circuit.

**Figure 5.**
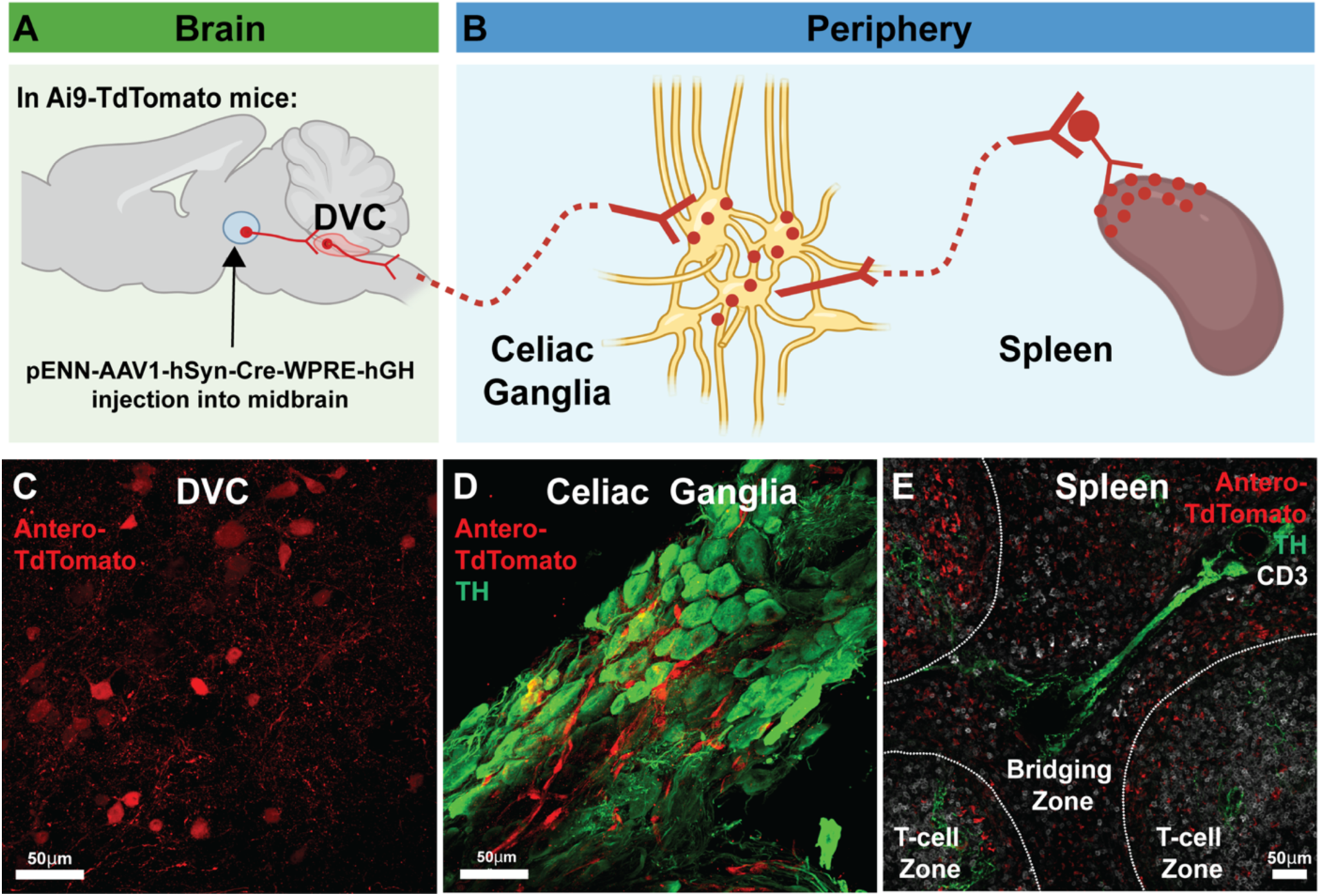
Anterograde multi-transsynaptic tracing reveals midbrain-to-spleen connectivity via the celiac ganglion. **Panel A:** Sagittal view of the mouse brain, showing multi-transsynaptic site of midbrain, where Ai9-Flox-stop-TdTomato mice were injected with pENN-AAV1-hSyn-Cre-WPRE-hGH. **Panel B:** Diagram showing where, after three weeks of viral expression, the DVC, celiac ganglia and spleen were assessed via confocal microscopy for expression of TdTomato. **Panel C:** DVC neurons are positive for TdTomato, indicating midbrain-to-DVC projections, in a manner that is consistent with our prior findings. **Panels D-E:** Assessment of celiac ganglia and spleen show presence of TdTomato^+^ neurons in both organs, indicating both organs received multi-transsynaptic projections from midbrain dopamine neurons. Taken together with our earlier neural circuit tracing data, these findings indicate that multi-synaptic projections exist from midbrain dopamine neurons, via DVC and celiac ganglia, to the spleen. *Data representative of n=4 independent experiments*.

### Chemogenetic stimulation of midbrain dopamine neurons increases cFos expression in the DVC and celiac ganglion

Thus far, we have shown: 1) that midbrain dopamine neurons mono-synaptically project to and release dopamine in the DVC; 2) that DVC neurons can respond to dopaminergic input, and 3) that multi-synaptic projections exist from the midbrain to the spleen via a midbrain-DVC-celiac ganglion-spleen circuit. This led us to ask whether activation of midbrain dopamine neurons increase neuronal activity along the midbrain-DVC-celiac ganglion axis. Immunostaining for cFos expression, an immediate-early gene induced by neuronal activity that reaches peak expression within 90 minutes after neuronal activation^67-70^, is a reliable measure of increased neuronal activity along the midbrain-DVC-celiac ganglion axis. Therefore, we next quantified cFos expression in the DVC and celiac ganglion as a proxy indication of neuronal activity. We injected AAV2-hSyn-DIO-hM3D(Gq)-mCherry or AAV2-hSyn-DIO-mCherry (control vector) into the midbrain of DAT-Cre mice. The same animals also received either a cervical vagotomy, or sham surgery targeting the cervical region. Four weeks post-injection, the mice received daily intraperitoneal injections of CNO (1 mg/kg) for 14 days to activate the Gq-DREADD-expressing midbrain dopamine neurons (Figure 6A). The choice of 1 mg/kg CNO was guided by prior work reporting that this dose does not elicit reward-seeking behavior or toxicity^71^. In addition, to minimize ligand-related confounds, all animals, both Gq-DREADD and empty-vector controls, received the same CNO regimen throughout the experiment. Thus, any direct or metabolite-mediated effects of CNO should be similar across all experimental and control groups.

**Figure 6.**
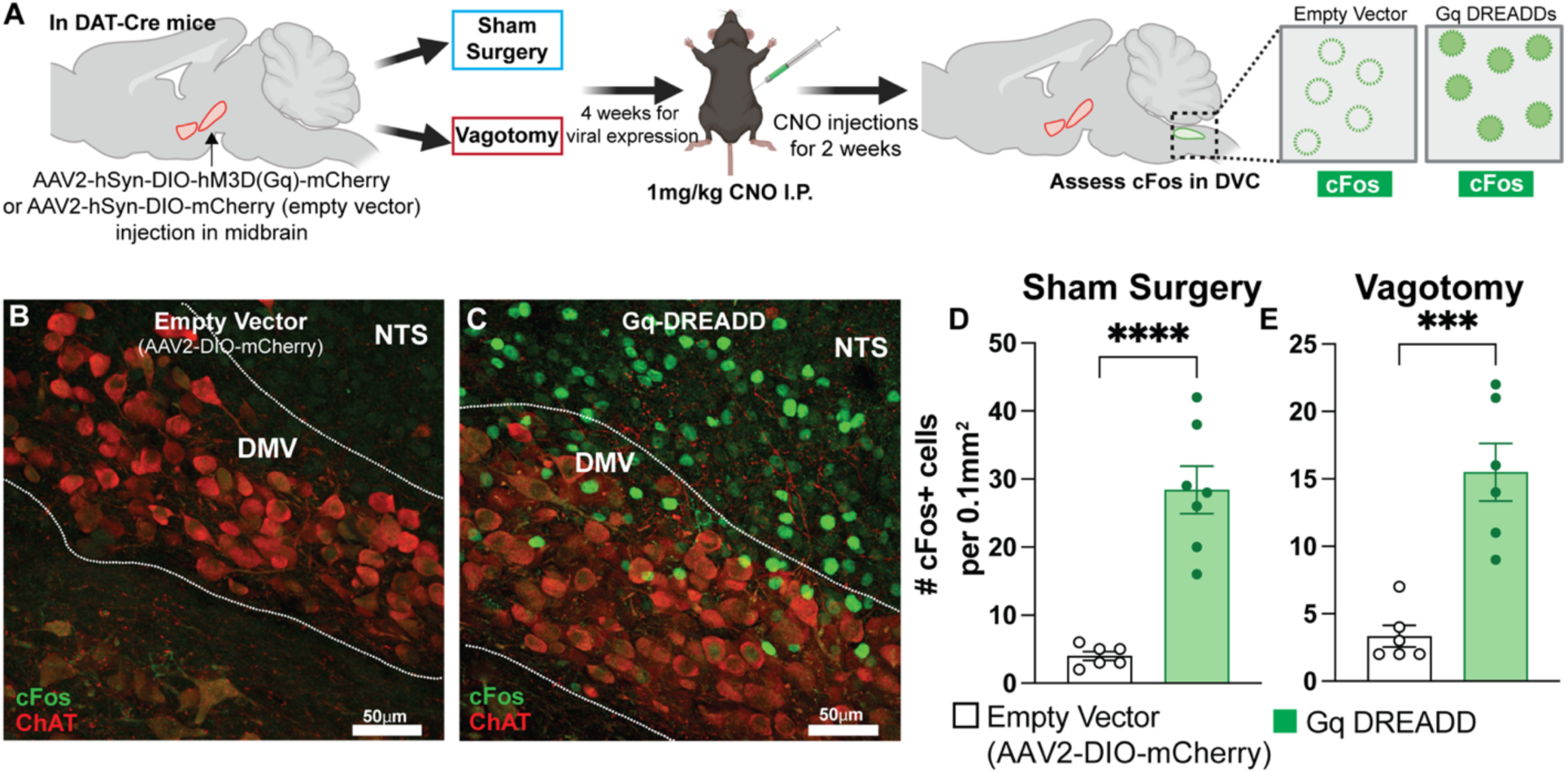
Stimulation with Gq-DREADD of midbrain dopamine neurons increases cFos expression in the DVC, indicating that dopamine neuron activation can modulate DVC activity. **Panel A:** Experiment timeline, showing (left) site of AAV2-hSyn-DIO-hM3D(Gq)-mCherry or empty vector (AAV2-hSyn-DIO-mCherry) injections into the midbrain DAT-Cre mice, (middle-left) four-week expression interval, (middle-right) daily I.P. injections of clozapine-*N*-oxide (1mg/kg), and (right) cFos immunolocalization in DVC as a proxy measure of neuronal activation. **Panel B:** Minimal/basal cFos immunostaining in the DVC of mice receiving empty vector (AAV2-hSyn-DIO-mCherry). **Panel C:** Robust cFos immunostaining in midbrain of mice injected with AAV encoded with Cre-dependent Gq-DREADD and treated with CNO. **Panel D:** Compared to empty vector control, mice receiving AAV2-hSyn-DIO-hM3D(Gq)-mCherry show significantly increased number of cFos+ cells in the DVC. **Panel E**: Following cervical vagotomy, animals that received AAV2-hSyn-DIO-hM3D(Gq)-mCherry continue to show increased DVC cFos expression compared to empty vector control. (*Data representative of n=6-7 independent biological replicates; ***p<0.005, ****p<0.0005, two-tailed unpaired Student’s T-test*)

Locomotor activity, a reliable readout of dopamine neuron activation, was assessed at baseline and following CNO injections on Days 1, 7, and 13 (Supplementary Figure 6 and 7; Supplemental Video 1 and 2). Mice injected with AAV2-hSyn-DIO-hM3D(Gq)-mCherry increased locomotor activity in response to CNO, confirming activation of midbrain dopamine neurons (Supplementary Figure 6B & 6C). Consistent with our hypothesis, we observed a robust increase in cFos expression in the DVC, particularly within the DMV and NTS subregions (Figure 6); increased DVC cFos expression was present in both sham surgery (Figure 6D) and vagotomized mice (Figure 6E). This effect was absent in the empty-vector control groups (AAV2-hSyn-DIO-mCherry) (Figure 6B-E). cFos expression in the DVC was detected in both ChAT⁺ and ChAT⁻ neurons, consistent with the heterogeneous neuronal populations in this region and its role as a relay hub that integrates and bidirectionally transmits information between the CNS and peripheral organs^72,73^ ^74,75^. We did not detect cFos expression in any of amygdala’s subregions that are known to receive inputs from, and send projections to, the midbrain and DVC (Supplemental Figure 8). The observation that midbrain dopamine neuron activation increased neuronal activity across multiple DVC subregions, including cholinergic neurons in the DMV and catecholaminergic neurons in the NTS agrees with prior reports^72,73^ ^74,75^, supporting the overall interpretation that activation of midbrain dopamine neurons influences neuronal activity along the midbrain-DVC axis, as evidenced by increases in cFos expression.

**Figure 7.**
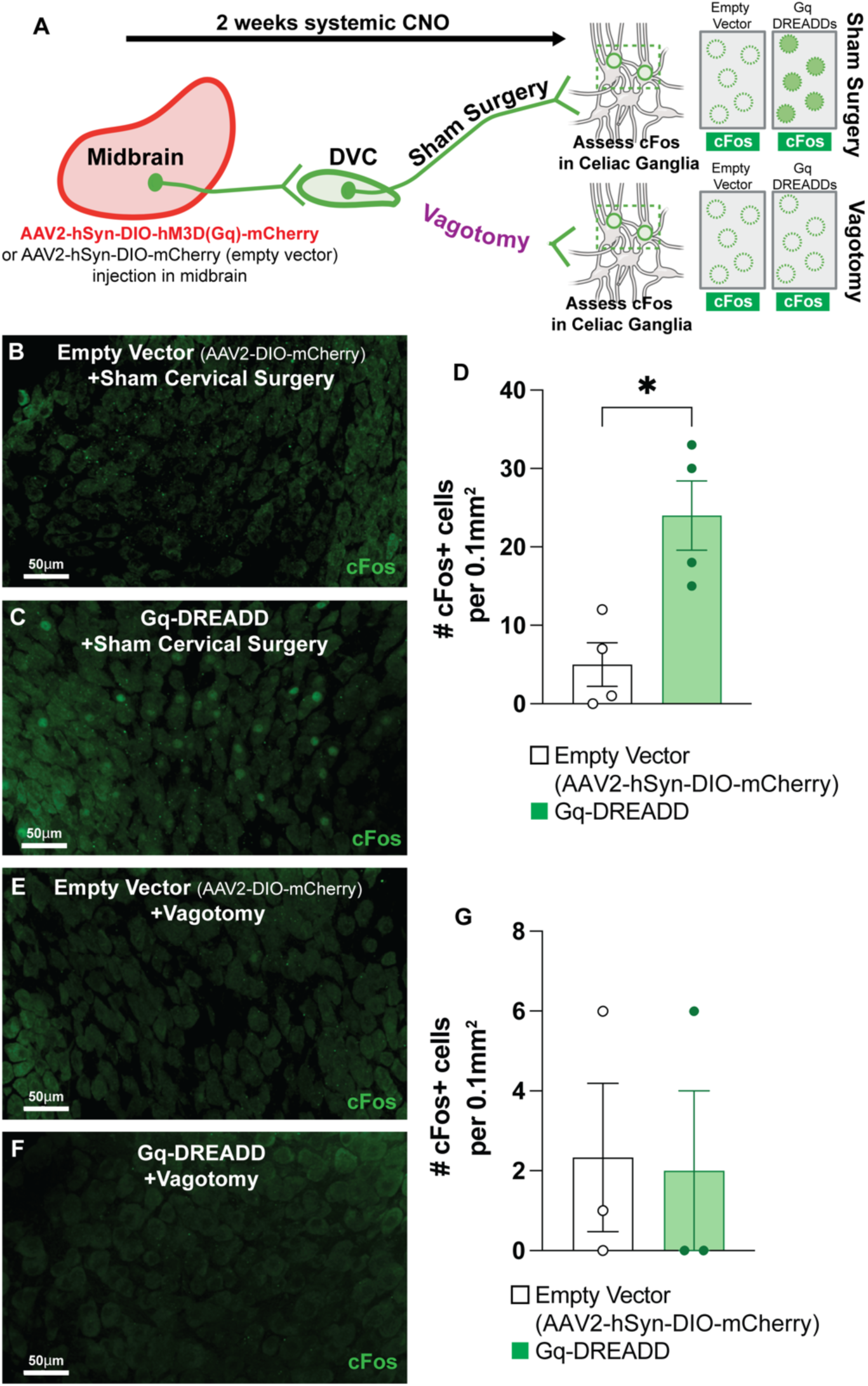
Gq-DREADD stimulation of midbrain dopamine neurons increases cFos expression in celiac ganglia, indicating that dopamine neuron activity can modulate celiac ganglia activity via midbrain-DVC-celiac ganglion pathway. Cervical vagotomy abolishes increased cFos expression in the celiac ganglia. **Panel A**: Cre-dependent excitatory Gq-DREADDs or empty vector control (AAV2-hSyn-DIO-mCherry) was injected in the midbrain of DAT-Cre mice. Following, mice received either a sham cervical surgery (top) or cervical vagotomy (bottom). After 4 weeks to allow viral expression, mice received 14 days of daily CNO (1mg/kg, I.P.). cFos immunoreactivity was examined as a proxy measure for increased celiac ganglion neuronal activity due to midbrain dopamine neuron activation. **Panel B-C**: Compared to empty vector control, animals that received Gq-DREADDs and I.P. CNO show increased celiac ganglion cFos expression. **Panel D**: Mice receiving Gq-DREADDs show significantly increased number of cFos+ cells in the celiac ganglion. **Panel E-F**: In mice that received cervical vagotomy, Gq-DREADD mediated stimulation of midbrain dopamine neurons does not significantly increase the number of cFos+ cells in the celiac ganglion. **Panel G**: Mice that received cervical vagotomy do not show increased (*Data representative of n=3-4 independent biological replicates; *p<0.05, two-tailed unpaired Student’s T-test*)

To further investigate downstream effects of midbrain dopamine neuron activation, we assessed cFos expression in the celiac ganglia of mice injected with Gq-DREADDs (excitatory DREADD) or control vector, and either cervical vagotomy or sham cervical surgery, as described above (Figure 7A). Following 14 days of CNO injection (1mg/kg, *i.p*.), intact celiac ganglia were dissected, whole-mounted, post-fixed, and immunostained. Consistent with midbrain-DVC-celiac ganglion circuitry, we observed increased neuronal activity, as evidenced by enhanced cFos expression in Gq-DREADD-injected mice, compared to the empty vector control (AAV2-hSyn-DIO-mCherry) (Figure 7B & 7C). Quantification of cells with robust cFos expression showed that, compared to mice that received the control vector which showed only basal levels of cFos expression, Gq-DREADD-treated mice receiving CNO revealed a marked increase in cFos+ cells (Figure 7D).

Since the celiac ganglion receives diverse inputs^53,59,76^ and innervates multiple peripheral organs^53,64,77,78^, we considered the possibility that the increased celiac ganglion activity observed after chemogenetic activation of dopamine neurons could arise form a parallel or indirect pathway. A straightforward way to test this was to disrupt the proposed DVC–celiac ganglion circuit by transecting the vagus nerve. Thus we tested whether increased celiac ganglion activity (in the context of midbrain dopamine neuron activation) persists following cervical vagotomy. An additional cohort of DAT-Cre mice received midbrain injections of either Gq DREADDs or empty vector control, followed by a sham surgery or a cervical vagotomy (Figure 7A). After 4 weeks to allow vial expression and surgical recovery, animals received systemic CNO as described above. Locomotor activity was assessed to verify dopamine neuron activation in both sham surgery and vagotomized mice (Supplemental Figures 6 and 7; Supplemental Video 1 and 2). Intact celiac ganglia were dissected, whole mounted, post-fixed and immunostained. Consistent with our hypothesis, animals that received both midbrain Gq-DREADD AAV and cervical vagotomy exhibit no change in celiac ganglion cFos expression relative to empty vector controls that received cervical vagotomy (Figure 7E-G). These results strongly support the interpretation that midbrain dopamine neuron activity functionally regulates splenic immunity via a midbrain-DVC-celiac ganglion-spleen circuitry. Together, these findings demonstrate that midbrain dopamine neuron activation profoundly influences neuronal activity along the midbrain-DVC axis and downstream autonomic circuits, highlighting a functional role in regulating peripheral immune responses.

### Chemogenetic stimulation of midbrain dopamine neurons reduces spleen weight and naïve CD4^+^ T-cell populations without affecting total cell numbers. Vagotomy abolishes changes in spleen weight and changes in naïve CD4^+^ T-cells

As a key immune organ, the spleen plays a central role in orchestrating both innate and adaptive immune responses, responding to pathogens, immune activation, and even cancer^79-86^. Disruptions in splenic immune cell populations or function due to altered CNS dopamine neurotransmission could have profound immunological consequences. Such behavior is particularly relevant in diseases characterized by dysregulated dopamine neuron activity and existing peripheral immune disturbances, potentially influenced by CNS dopamine signaling. For instance, Parkinson’s disease (PD), conventionally defined by the loss of midbrain dopamine neurons and motor deficits, also exhibits peripheral immune manifestations, including changes in the phenotype of immune cell populations, gut dysfunction, and systemic inflammation^3-5,87^, with PD patients exhibiting impaired CD4^+^ T-cell migration, accompanied by an increase total CD4^+^ T-cell populations^87-89^. Conversely, psychostimulant use, which elevates brain dopamine levels, leads to opposing T-cell responses, including a reduction in total T-cells, and an increased propensity towards inflammatory T-cell phenotypes^90-94^. These observations suggest that changes in CNS dopamine transmission, whether through loss, as in PD, or through augmentation, as seen with psychostimulants, can influence immune function^87-89,90-94^. The interplay between midbrain dopamine neuron activity and peripheral immune responses underscores a relationship where CNS signaling impacts systemic immunity^6,95-98^. To probe whether the midbrain-DVC-celiac ganglion-spleen circuit modulates splenic immunity, we assessed splenic immune cell populations following the activation of midbrain dopamine neurons. The proposed midbrain-DVC-celiac ganglion-spleen circuit relies on vagal projections from DVC to spleen via the celiac ganglion^24,25^; therefore, we reasoned that the effects of midbrain dopamine neuron activation on splenic immunity would be abolished if the vagus nerve were transected. Previous work has shown that cervical, but not subdiaphragmatic, vagotomy preserves connections between the celiac ganglion and spleen while disrupting vagal efferents to the celiac ganglion^99,100^.

Therefore, we employed a four-pronged experimental design, in which DAT-cre mice received an intracranial AAV injection targeting the midbrain to express either Cre-dependent Gq-DREADDs or a control vector expressing a fluorescent reporter, followed by either a left cervical vagus transection (vagotomy) or a sham surgery. After four weeks to allow viral expression, we administered CNO daily (1 mg/kg, I.P.) for 14 days to stimulate DAT^+^ midbrain neurons (Figure 8A, Figure 9A). Locomotor activity was assessed at baseline and following CNO injections on Days 1, 7, and 13 to verify chemogenetic stimulation of dopamine neurons (Supplementary Figure 6 and 7; Supplemental Video 1 and 2). Chemogenetic stimulation of dopamine neurons did not alter body weight in either sham surgery or vagotomy groups (Figure 8B and Figure 9B),

**Figure 8.**
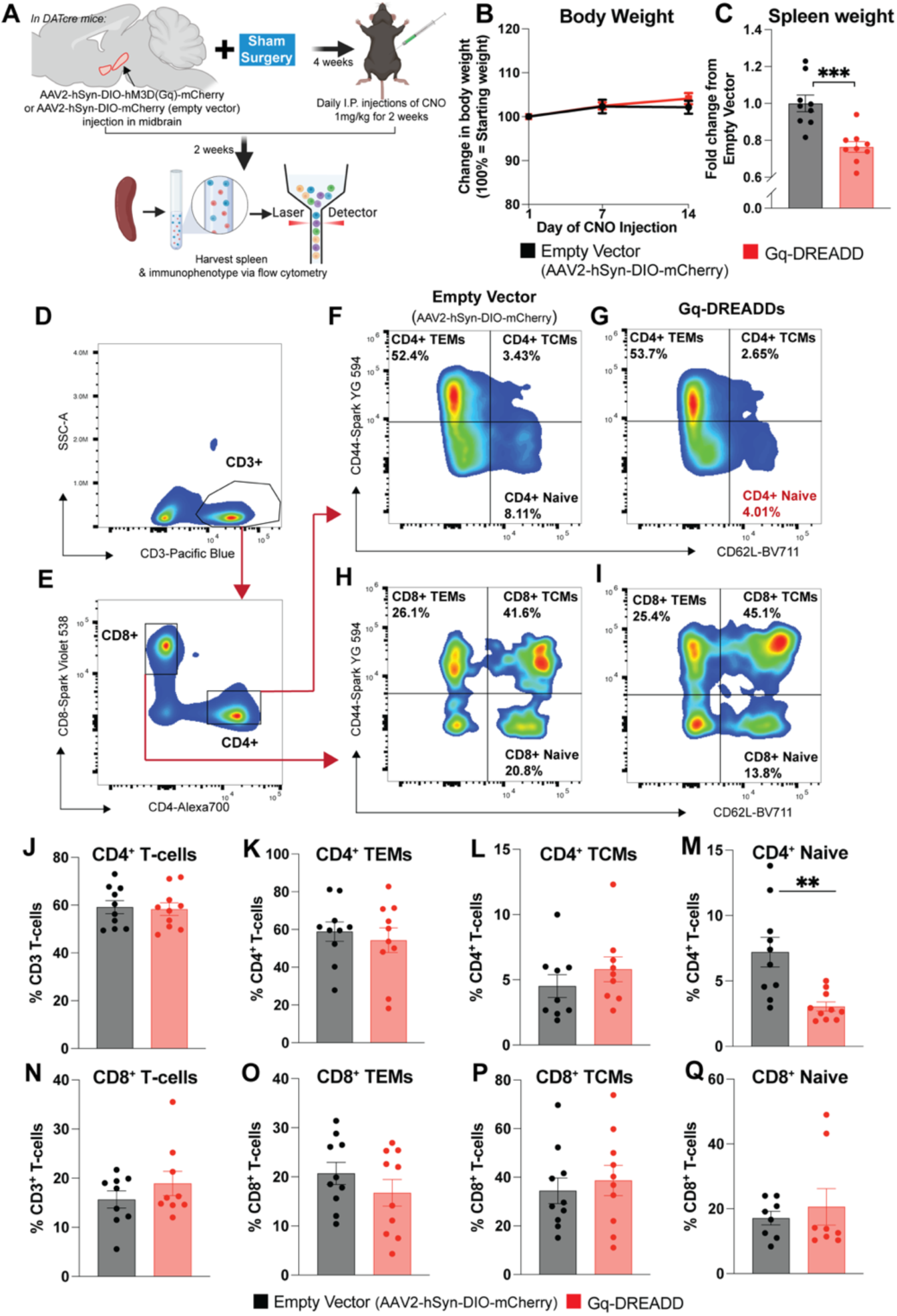
Spleen weight and flow cytometric analysis, showing that stimulation of midbrain dopamine neurons by Gq-DREADD results in reduced spleen weight and reduced naïve CD4^+^ T-cell numbers in the spleen without effect on CD8^+^ T-cell populations. **Panel A:** Experiment timeline, showing site of AAV2-hSyn-DIO-hM3D(Gq)-mCherry or empty vector (AAV2-hSyn-DIO-mCherry) injections into the midbrain DAT-Cre mice, followed by sham surgery targeting the cervical region. After four-weeks for viral expression, followed by daily I.P. injections of clozapine-*N*-oxide (1mg/kg) for two weeks, harvesting spleen for weighing cellular dissociation, and (right) flow cytometry using antibodies against T-cell subtypes. **Panel B:** Animal body weight was monitored throughout CNO administration; mice that received Gq-DREADDs vs empty vector control did not exhibit decreases in body weight. **Panel C**: Reduced spleen weight (adjusted for body weight) in mice receiving midbrain Gq-DREADD and systemic CNO, as compared to empty vector control (p<0.05). **Panels D-E**: Single live cells, free of debris, were gated for CD3 expression, a pan-T-cell marker, and then gated for expression of CD4 and CD8. **Panels F-I:** CD4^+^ and CD8^+^ T-cells were analyzed by expression of CD44 and/or CD62L to assess populations of T-effector memory cells (TEMs), T-central memory cells (TCMs) or naïve T-cells. **Panels H-M:** With no change in total CD4+ T-cell populations, compared to empty vector control, animals that received Gq-DREADDs in the midbrain show significantly reduced naïve CD4+ T-cells, marked by expression of CD62L and lacking expression of CD44. CD4^+^ T-effector memory cells and T-central memory cells were not significantly changed. **Panels N-Q:** CD8^+^ T-cell subsets are not significantly changed by Gq-DREADD-mediated midbrain dopamine neuron activation. *(n=8-10 independent biological replicates. Panel B: p>0.05, 2-way ANOVA, N.S. Panels J-Q: *p<0.05, two-tailed unpaired Student’s T-test)*

**Figure 9.**
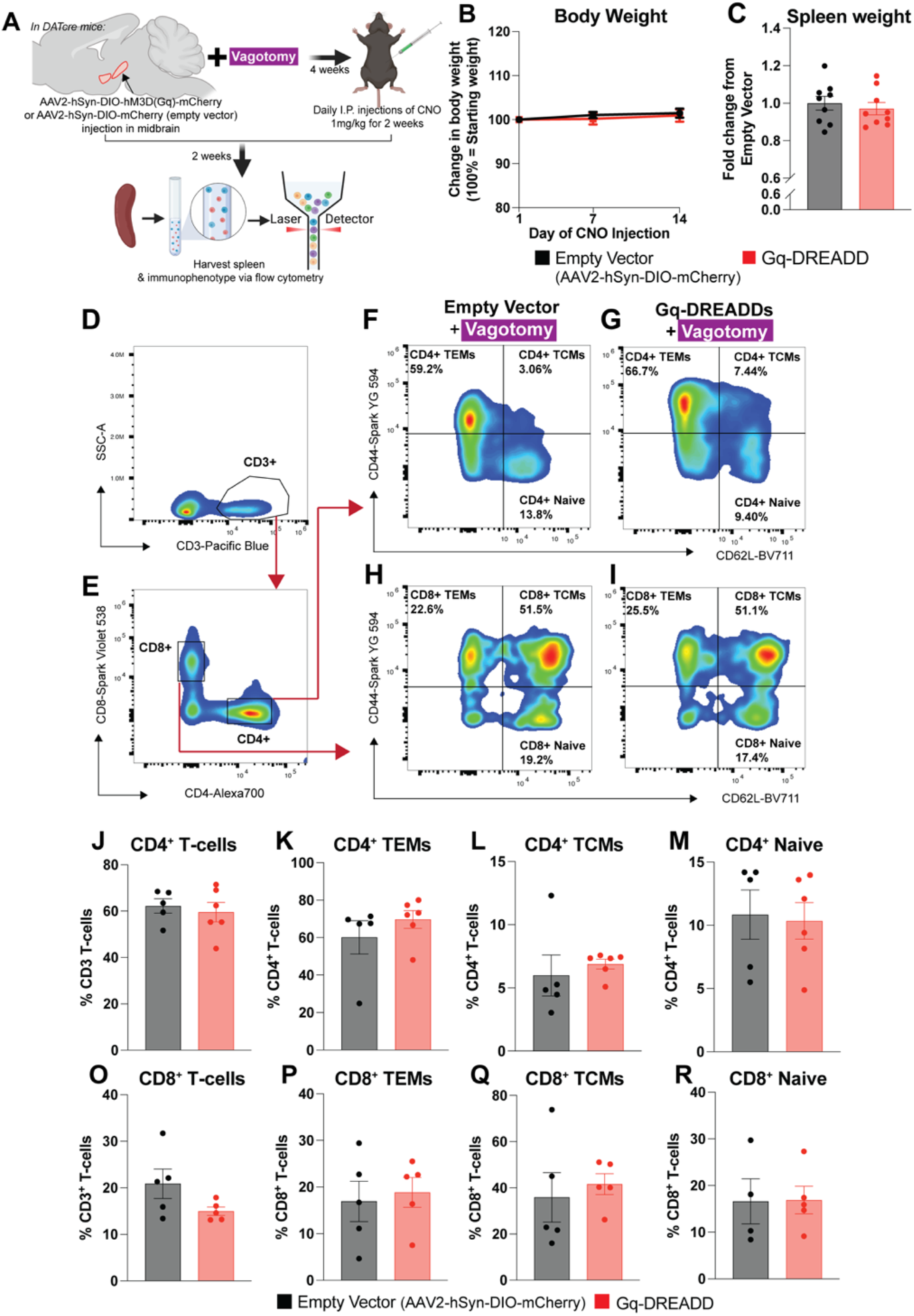
Following vagotomy, effect on spleen weight and splenic immune cells is abolished despite activation of midbrain dopamine neurons, indicating a specific pathway from midbrain to spleen via vagal afferents. **Panel A:** Experimental timeline showing site of AAV2-hSyn-DIO-hM3D(Gq)-mCherry or empty vector (AAV2-hSyn-DIO-mCherry) injections into the midbrain DAT-Cre mice, followed by cervical surgery transecting the vagus nerve (cervical vagotomy). After four-weeks for viral expression, followed by daily I.P. injections of clozapine-*N*-oxide (1mg/kg) for two weeks, harvesting spleen for weighing and cellular dissociation, and analysis flow cytometry using antibodies against T-cell subtypes. **Panel B**: Animal body weight was monitored throughout CNO administration; mice that received Gq-DREADDs vs empty vector control did not exhibit decreases in body weight. **Panel C**: In contrast to intact animals receiving midbrain Gq-DREADDs, vagotomized animals that received Gq-DREADDs or Empty Vector Control did not exhibit significantly different spleen weights adjusted for body weight. **Panels D-E**: Single live cells, free of debris, were gated for CD3 expression, a pan-T-cell marker, and then gated for expression of CD4 and CD8. **Panels F-I:** In vagotomized animals that received either empty vector control or Gq-DREADDs, CD4^+^ and CD8^+^ T-cells were analyzed by expression of CD44 and/or CD62L to assess populations of T-effector memory cells (TEMs), T-central memory cells (TCMs) or naïve T-cells. **Panel J-R**: Regardless of whether animals received empty vector control or Gq-DREADD AAV in the midbrain, cervical vagotomy abolishes the impact of midbrain dopamine neuron activation on splenic T-cell populations *(n=4-6 independent biological replicates. Panel B: p>0.05, 2-way ANOVA, N.S. Panels J-Q: p>0.05, two-tailed unpaired Student’s T-test, N.S.)*.

Firstly, examining the sham surgery group, we found that mice with chemogenetic stimulation of midbrain dopamine neurons showed a significant reduction in spleen weight relative to empty vector controls (Figure 8C), despite no change in body weight, reinforcing the idea that midbrain dopamine signaling influences splenic immune function. These data are consistent with previous studies in DAT-knockout (DAT-KO) mice, marked by elevated CNS dopamine, which is associated with spleen hypoplasia, and reduced spleen weight^98,101^. Next, in the sham surgery group, we performed flow cytometry to assess splenic adaptive and innate immune subsets. These analyses included T-cell populations, including CD4^+^, CD8^+^, effector memory, Central memory, and naïve T-cells (Figure 8C-H, Supplemental Figure 6); B-cell populations, including total B-cells, immature and maturing B-cells (Supplemental Figures 7 and 12); NK-cell populations, including total NK-cells, immature and mature NK-cells (Supplemental Figures 8 and 12); and splenic macrophages (Supplemental Figures 10 and 14). While total numbers of CD4^+^ T-cells, effector memory T-cells (CD4^+^ CD44^+^ CD62L^-^) and central memory T-cells (CD4^+^ CD44^+^ CD62L^+^) were unchanged between groups (Figure 8 F-G; J-L), naïve CD4^+^ T-cells (CD4^+^ CD44^-^ CD62L^+^) were significantly reduced in mice that received chemogenetic stimulation of midbrain dopamine neurons, compared to empty vector control (Figure 8M). No significant changes were observed in CD8^+^ T-cell populations (Figure 8 H-I; N-Q).

To determine the involvement of the vagus nerve in modulating splenic immunity in the proposed midbrain-DVC-celiac ganglion-spleen when dopamine neurons are chemogenetically activated, we examined the groups that received midbrain injection of Gq-DREADD AAV or empty vector control followed by vagotomy (Figure 9). In contrast to the sham surgery groups, in vagotomized mice, chemogenetic stimulation of midbrain dopamine neurons show no change in spleen weight compared to empty vector controls (Figure 9C). Immunophenotyping of splenic immune cells in vagotomized mice showed that total CD4+ T-cells, effector memory cells (CD4+ CD44^+^ CD62L^-^), and central memory cells are unchanged (CD4^+^ CD44^+^ CD62L^+^) (Figure 9J-L), similar to sham controls. Importantly, diverging from sham surgery groups, we found that vagotomy abolishes the reduction in naïve CD4^+^ T-cells (CD4^+^ CD44^-^ CD62L^+^) in mice with chemogenetic stimulation of midbrain dopamine neurons, compared to empty vector controls (Figure 9 F-G, Figure 9M). No changes were observed in CD8^+^ T-cell populations in both groups of vagotomized mice (Figure 9H-I, Figure 9N-Q). Finally, in both sham surgery and vagotomy groups, we observed no changes in B-cell subtypes (Supplementary Figure 9), NK-cell subtypes (Supplementary Figure 10) or splenic macrophages (Supplementary Figure 11).

Collectively, these findings suggest a brain-body circuit for specialized modulation of specific splenic immune cell populations by midbrain dopamine neurons, rather than global suppression, shedding light on mechanisms by which altered dopamine neurotransmission influences peripheral immune function. Importantly, our results align with previous studies demonstrating that top-down regulation of splenic T-cells can affect peripheral immune manifestations such as pain^96^.

Through the celiac ganglia, the spleen is innervated by both cholinergic^29-31^ and catecholaminergic fibers^32-35^, both of which can modulate spleen function^26^, immune cell populations, and, by extension, influence the peripheral immune landscape^102-104^. Therefore, dopaminergic projections from midbrain regulate DVC activity, that via the vagus nerve and celiac ganglion with downstream adrenergic and cholinergic fibers, could modulate splenic immunity^102-104^. This pathway likely underpins the immunophenotypic changes observed following midbrain dopamine neuron activation. These are consistent with the central finding of the current study that activation of midbrain dopamine neurons induces functional immunological changes in the spleen. It should be noted that the DVC contains multiple vagal nuclei upstream of the periphery and projects to numerous peripheral organs via the vagal efferent pathway. This raises the possibility that the changes in splenic immunity observed upon chemogenetic activation of midbrain dopamine neurons may also reflect alterations in the activity of other peripheral organs outside the proposed midbrain-DVC-celiac ganglion-spleen circuit^38,51,75,105^.

### Concluding remarks

This study provides strong evidence for a functional connection between the CNS and peripheral immunity through midbrain dopamine signaling. Our findings suggest that midbrain dopamine neuron activation not only affects splenic weight and T-cell populations but also underscores a broader role for dopamine signaling in regulating peripheral immunity. Disruptions in brain-body communication have been implicated in various diseases, including neurodegenerative disorders^3^, cardiovascular disorders^106-108^, and metabolic syndromes^109,110^. Numerous studies have shown that altered dopamine neurotransmission is linked to changes in peripheral immunity, such as during psychostimulant use and with degeneration of dopamine neurons^6,87,97,98,111-113^. However, prior evidence for the circuitry between midbrain dopamine neurons and the immune system was scant. In the present study, we now demonstrate the existence of a multi-synaptic brain-to-periphery pathway, linking midbrain dopamine neurons to the DVC and, via the celiac ganglia, to the spleen^26^. Vagal fibers are known to innervate the celiac ganglion^25,99,100,114^; in turn, projections from the celiac ganglion innervate the spleen^114^. Given the evidence linking the vagus to the spleen, prior studies have focused on the cholinergic anti-inflammatory pathway that can modulate inflammation via vagal transmission^26-28^, further supporting the importance these of brain-body connection.

While our findings demonstrate that midbrain dopamine neuron activity, acting through a midbrain–DVC–celiac ganglion pathway, modulates splenic immunity, our data do not exclude contributions from parallel routes within the vagal–spleen axis. Indeed, the vagus nerve sends and receives projections from the gastrointestinal tract^36,115-117^, and gut–spleen communication via autonomic and immune pathways are well documented^104,118,119^. These observations raise the possibility that a midbrain–DVC–gut–spleen circuit could contribute to midbrain dopamine neuron–dependent regulation of peripheral immunity. Supporting this idea, Chang et al. (2024) identified a stress-sensitive circuit in which vagal projections regulate gut microbiome composition, and altered gut microbiota influence both splenic immunity and brain function^116,120-122^. Thus, a brain–gut–spleen axis represents a plausible complementary pathway for dopamine neuron modulation of peripheral immunity. Finally, although the present study focuses on the efferent limb of the midbrain–DVC–celiac ganglion–spleen pathway, our data do not preclude afferent signaling from the periphery back to the CNS and the existence of other reciprocal, bidirectional circuits between these compartments^123-125^.

In the present study, we expand previous findings demonstrating that midbrain-originating dopamine signaling emerges as a key modulator of peripheral immunity, with the observed reduction in spleen weight and alterations in T-cell populations, particularly the decrease in naïve CD4^+^ T-cells, following chemogenetic activation of midbrain dopamine neurons. These results emphasize the influence of midbrain dopamine neurons on adaptive immune functions, with broad implications for conditions where immune dysregulation and chronic inflammation are central to disease manifestation and progression. Furthermore, dopamine-driven modulation of splenic immune cell populations suggests that midbrain dopamine neuron activity may extend regulatory effects to immune responses throughout the body. A multitude of diseases, from autoimmunity to cancer, are treated using immunomodulatory drugs, modulating the immune response as a treatment method. Given that dopamine neuron activity affects splenic T-cell populations, we pondered whether peripheral immunomodulation may also change CNS dopamine neurotransmission, via the same circuit. It is well known that during infection and inflammation, patients experience a lack of motivation, social withdrawal, and anhedonia^12,126-128^. Therefore, it is reasonable to postulate that peripheral immunomodulation, via drugs that target the immune system, may function in a feedback/feedforward loop to modulate CNS dopamine neurotransmission. Future studies employing clinically relevant immunomodulatory drugs, or experimental methods such as chemogenetic manipulation of the spleen, will help clarify this compelling question.

Finally, there remains the fundamental question: why have mammals evolved a biological circuit that links midbrain dopamine neurons to the peripheral immune system? Dopamine neurotransmission is fundamentally involved in socially motivated behaviors, such as social bonding^129-133^. Such behaviors are conserved among numerous species. Indeed, in contexts of anhedonia, such as during depression, peripheral inflammation and immune dysfunction is increased^134,135^. In this respect, midbrain dopamine regulation of the peripheral immune system may facilitate immunity through socialization and bonding reward mechanisms. This idea is supported by literature showing that activation of midbrain dopamine neurons helps to resolve infection^95^, to improve anti-tumor immunity^96^, and to enhance stress resilience in individuals experiencing chronic stress^136,137^, emphasizing the function of dopaminergic signaling in peripheral immunity along the midbrain-spleen axis. Our study also highlights a critical role for midbrain dopamine neurons beyond CNS neurotransmission, positioning them as an important regulator of peripheral immunity. Such considerations strengthen the concept of brain-body communication, linking neuroimmune interactions to the onset and progression of systemic diseases and offering promising leads for therapeutic interventions that target these circuits.

### Limitations of the methodology and systems used in this study

While our approach yielded significant findings, several methodological limitations warrant consideration. First, the viral constructs employed in our study reliably transduced the entire dopaminergic midbrain, encompassing both the substantia nigra pars compacta (SNc) and the ventral tegmental area (VTA), precluding efforts to distinguish the specific roles of SNc versus VTA neurons in the regulation of the peripheral immune system. Future studies utilizing more refined viral constructs or intersectional genetics will enable dissection these distinct neuronal populations and likely would uncover even more robust outcomes. Second, our data suggest that midbrain dopamine neurons send projections throughout the DVC, including AP, NTS, and DMV, thus preventing us from determining whether a specific nucleus or neuronal subtype within the DVC contributes predominantly to splenic immunoregulation by midbrain dopamine signaling. Future studies employing targeted inhibition of individual DVC nuclei, during dopamine neuron activation will help delineate the specific roles of each nucleus and individual cell types in splenic immune modulation. Third, literature suggests that unilateral vagus nerve stimulation may induce more pronounced changes in the peripheral immune system^138,139^; however, our bilateral approach does not allow us to assess whether the left or right vagal pathway exerts greater influence on midbrain dopamine neuron regulation of splenic immunity. Addressing this limitation in future studies will involve investigating the laterality of midbrain-spleen immunoregulation, which could provide deeper insights into the lateral specificity of this pathway. Such findings could have important implications for experimental designs and therapeutic applications based on these mechanisms. While we have taken a significant step toward understanding the role of midbrain dopamine signaling in regulating peripheral immunity, addressing these constraints should provide a fuller perspective on the neuroimmunological implications of the midbrain-DVC-celiac ganglion-spleen pathway.

## Materials and Methods

### Animals

We used the following strains of mice, both males and females, on a C57BL/6 background: B6.Cg-*Gt(ROSA)26Sor^tm9(CAG-tdTomato)Hze^*/J (Jax strain #007909)^140^, *Slc6a3^tm1(cre)Xz^*/J (Jax strain #020080), B6.Cg-Tg(RP23-268L19-EGFP)2Mik/J (Jax strain#007902). Adult male and female mice were between 8-16 weeks of age, on a 12-hour day/night cycle with ad libitum access to food and water. Animals were bred in-house at the University of Florida or at the University of Texas at San Antonio, following approved procedures by the University of Florida IACUC (IACUC 202400000184) or by the University of Texas at San Antonio IACUC (IACUC MU096).

### Stereotactic surgeries and viral injections

Mice were anesthetized using gas anesthetic (3% isoflurane mixed with oxygen). After head shaving, mice were head-fixed in a stereotactic frame (Kopf Instruments). Following sterilization of the surgical site, an incision was made, skull exposed and leveled from bregma to lambda, and burr holes drilled at the surgical coordinates (Bregma: +0.4ML, -3.4AP, -4.4DV). Viruses were injected via a pulled glass capillary at a rate of 5nL every 5 seconds, until the injection was complete (Drummond Nanoject III). The injector was left in place for 5 minutes to allow spread prior to retraction. Scalp skin was sutured using 3 interrupted sutures (5.0 polypropylene sutures). The virus was allowed to express for several weeks prior to experiments as indicated throughout the Results. Viral vectors are provided in **Table 1**.

**Table 1:**
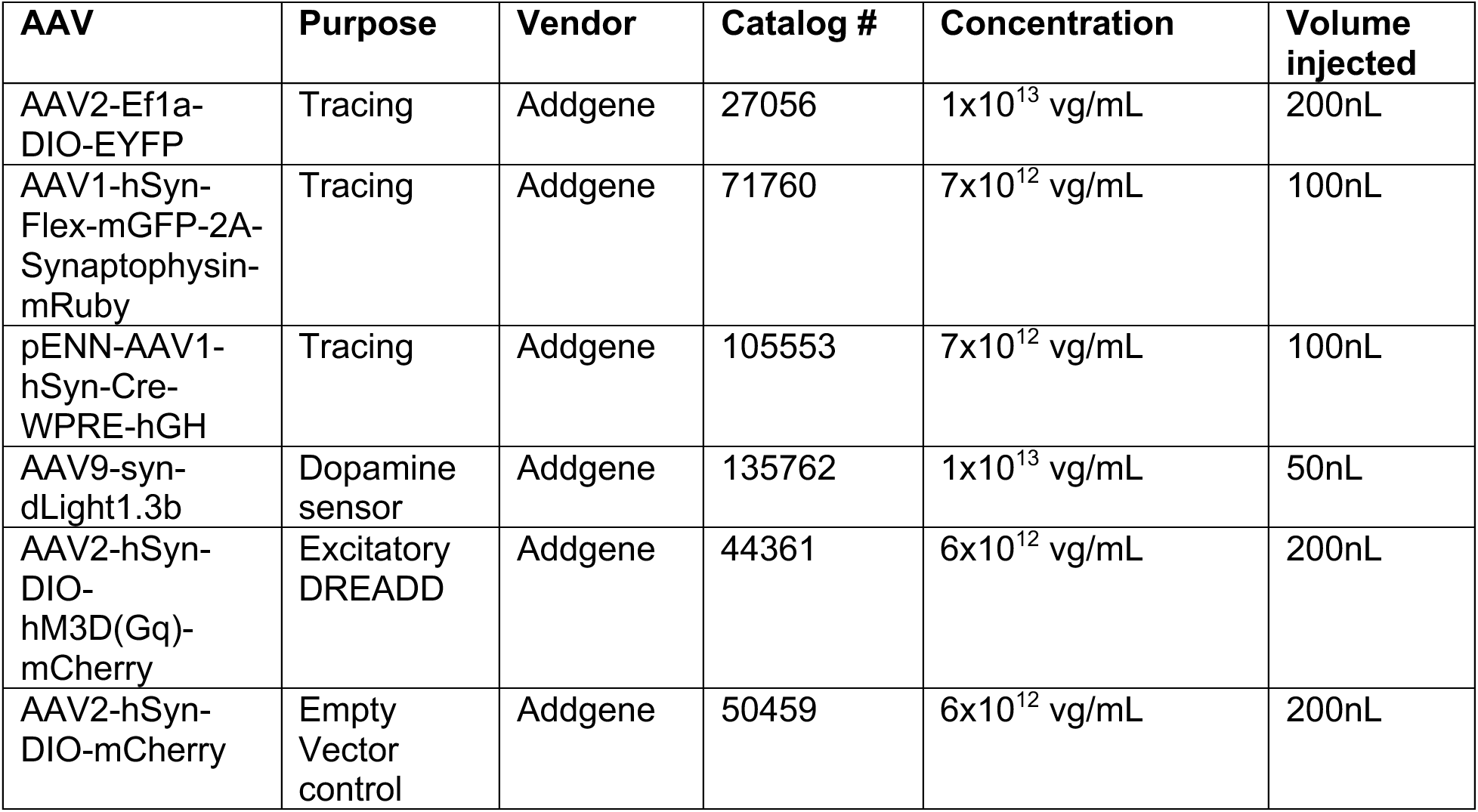
Adeno-associated viruses.

### Vagotomy or sham procedure

Mice were placed in a supine position on a sterile drape, maintained on isoflurane at 0.8-1.2%. Following sterilization of the neck surgical site, a vertical incision was made 3mm caudal to the sternal notch. The inferior border of the thyroid tissue, a distinct V-shaped line at the cervico-sternal junction, was identified. By blunt dissection, submaxillary salivary glands were exposed and separated through the midline fascial plane to expose the trachea. The incision was kept open using a micro-retractor. The left cervical vagus nerve, located within a neurovascular bundle, was readily identified by the pulsation of the carotid artery. The carotid sheath was isolated from the connective tissue using pickup forceps. The nerve was gently dissected from the sheath, taking care not to damage the vessels or tug on the nerve. Once isolated, a small section of sterile suture was used to suspend the nerve from surrounding tissue and severed using fine surgical scissors (vagotomy) or simply manipulated and then restored to the sheath (sham procedure). The incision site was cleaned and sutured with 3-5 interrupted sutures.

### Image analysis

Fluorescence intensity of EYFP+ processes in the DVC were calculated as total fluorescence intensity on background-corrected images in the channel corresponding to EYFP in NIS Elements software. Units are expressed as arbitrary fluorescence units per square micron. mRuby+ puncta were quantified using open-source software Cell Profiler, using an analysis pipeline adapted from Cell Profiler “Speckles Quantification”. cFos+ cells in the DVC and celiac ganglion were assessed by setting a threshold for cFos intensity of greater than or equal to 61% of the fluorescence intensity of the nuclear counterstain (DAPI). Cells which were positive for cFos were manually counted and are presented as number of cFos+ cells per square micron.

### Electrophysiology

As described in previous studies^141-147^, after 3-4 weeks from the viral injection, mice were anesthetized with isoflurane and decapitated. Coronal slices (300 μm) containing the area of interest were obtained on a vibratome (VT1200S, Leica) in a chilled cutting solution containing 100 mM choline chloride, 25 mM NaHCO_3_, 25 mM D-glucose, 11.6 mM sodium ascorbate, 7 mM MgSO_4_ 3.1 mM sodium pyruvate, 2.5 mM KCl, 1.25 mM NaH_2_PO_4_, 0.5 mM CaCl_2_. Slices were then incubated in oxygenated artificial cerebrospinal fluid (ACSF) in a submerged chamber at 35–37°C for 30 min and then at room temperature (21–25°C), until recordings were performed. ACSF contained 126 mM NaCl, 26 mM NaHCO_3_, 10 mM D-glucose, 2.5 mM KCl, 2 mM CaCl_2_, 1.25 mM NaH_2_PO_4_, 1 mM MgCl_2_; osmolarity was ∼290 Osm/L.

Whole-cell recordings were performed in 31–33°C ACSF. Thin-walled borosilicate glass pipettes (Warner Instruments) were pulled on a vertical pipette puller (PC-10, Narishige), resulting in a typical range of 3–5 MΩ resistance. Intrinsic properties were recorded in the current-clamp configuration using a potassium-based intracellular solution which contained 120 mM potassium gluconate, 20 mM KCl, 10 mM HEPES, 10 mM phosphocreatine, 4 mM ATP, 0.3 mM GTP, 0.2 mM EGTA, and 0.3–0.5% biocytin (mg/mL). Signals were sampled at 10 kHz and filtered (lowpass filter) at 4 kHz, in the presence of excitation and inhibition blockers: CPP (5 μM; Tocris Bioscience), NBQX (10 μM; Abcam), and gabazine (25 μM; Abcam). Hardware control and data acquisition were performed with the Matlab-based Ephus package (Suter et al., 2010). The resting membrane potential (Vm) was calculated in current clamp mode (I = 0) immediately after breaking in. Series (R_s_) and input resistance (R_in_) were calculated in voltage-clamp mode (V_hold_ = −70 mV) by giving a −5 mV step, which resulted in transient current responses. R_s_ was determined by dividing the voltage step amplitude by the peak of the capacitive current generated by the voltage step. The difference between baseline and steady-state hyperpolarized current (ΔI) was used to calculate Rin using the following formula: R_in_ = −5 mV/ΔI − R_s_. Subthreshold and suprathreshold membrane responses in current-clamp configuration were elicited by injecting −100 to +500 pA in 50-pA increments at V_hold_ = −70 mV. The first resulting AP at rheobase (the minimal current of infinite duration - experimentally limited to 1 s, as required to generate an AP, from here on current threshold, or I threshold) was analyzed for AP half-width, peak, height, Voltage threshold. The adaptation ratio was measured at the current step that gave the closest APs firing rate to 20 Hz, calculated dividing the second instantaneous frequency by the last. The f/I graphs were generated by calculating the firing frequency at each positive current step administered to the cell while holding it at -70 mV. Alpha synapse was calculated by measuring the peak of five consecutive 5 msec EPSP-like steps at 20Hz and normalizing on the amplitude of the first peak.

### *Ex vivo* dLight1.3b recording

#### Preparation of acute DVC slices

The mice were perfused with ice cold sucrose-sodium-replacement solution containing 250 mM sucrose, 26 mM NaHCO_3_, 2 mM KCl, 1.2 mM NaH_2_PO_4_, 11 mM dextrose, 7 mM MgCl_2_, and 0.5 mM CaCl_2_. Mice were decapitated and their brains were promptly removed. For slicing, the brain was cut rostrally to the cerebellum in the coronal plane to create a flat surface for attachment to the cutting stage. Coronal brainstem sections containing the DVC were sectioned at 250 µm (VT1200, Leica). The slices were transferred to a recovery chamber containing oxygenated ACSF at 34°C. The ACSF composition included: 126 mM NaCl, 2.5 mM KCl, 2 mM CaCl_2_, 26 mM NaHCO_3_, 1.25 mM NaH_2_PO_4_, 2 mM MgSO_4_, and 10 dextrose, equilibrated with 95% O_2_ and 5% CO_2_. The osmolarity for both the sucrose-sodium-replacement and ACSF was adjusted to 300-310 mOsm and the pH was set to 7.35-7.4. Sections remained in the recovery chamber for at least 30 minutes at 34°C until used for subsequent ex-vivo dLight1.3b recording. Sections containing the DVC were identified using the AP and hypoglossal nucleus as anatomical landmarks.

#### Recording

The slices were then transferred to a low-profile open diamond bath imaging chamber (RC-26GLP, Warner Instruments, Hamden, CT, USA) and perfused with ACSF at 36-37°C. Sections were imaged using a 40x water-immersion objective on an Eclipse FN1 upright microscope (Nikon Instruments, Inc., Melville, NY, USA), at 25-30fps with no delay between frames. The microscope was equipped with a Spectra X light engine (Lumencor, Beaverton, OR, USA) and a 470/24 nm solid-state illumination source. Images were captured with a Hamamatsu BT Fusion camera (Hamamatsu). Gq-DREADD receptors were activated via bath perfusion of 10 μM clozapine-N-oxide (CNO) in ACSF.

#### Signal bleach correction

Raw fluorescent intensity values were extracted using ImageJ. A photobleaching curve was generated by fitting the baseline florescent signal using a non-linear least regression with a biexponential model. Traces were then corrected and normalized using MATLAB. Codes used for bleach correction and normalization were generated by Leah T. Phan (co-author), and are available at: https://github.com/LTP08/Khosh_CommonCodes/blob/899d7403fb39b3d7d41978a11e6284de1787d55e/biexponential_photobleachcorrection_bulksignal.mlx

#### dF/F dlight1.3b signal calculation

Baseline values, the median value of the 30 second baseline recording (pre-drug) were subtracted from the peak signal 30 seconds following drug perfusion (∼1 minute after recording start). The resulting fluorescence value was divided by the baseline fluorescence value to yield a normalized dF/F value for comparison across experiments.

### Dissection of the sympathetic splanchnic-celiac-superior mesenteric ganglion complex (SCSMG)

Mice were deeply anesthetized with isoflurane, and transcardially perfused with PBS, followed by 4% paraformaldehyde (PFA). The sympathetic ganglion complex was dissected following Trevizan-Baú et al. (2024)^59^. Briefly, mice were positioned in dorsal recumbency, and the abdominal wall was opened (in the Linea Alba). With fine forceps and Noyes Iris scissors, the sympathetic ganglion complex was dissected and collected under a stereo microscope. The complex was then placed onto a Sylgard-coated dissection dish with PBS and pinned down on the dish through connective tissue and nerves, not through the ganglia. Then, after removing any additional connective tissue or fat, the sympathetic ganglion complex was post-fixed for 24 hours before using for immunostaining and later imaging.

### Brain tissue and spleen immunofluorescence and imaging

Animals were terminally anesthetized with isoflurane, and transcardiac perfused with 1x PBS followed by 4% PFA dissolved in PBS (pH 7.4). Brains were removed and post-fixed in 4% PFA for 24 hours at 4°C. Spleens were removed prior to perfusion and post-fixed in 4% PFA for 24 hours at 4°C. Thereafter, sections were cut at 40um thickness at room temperature on a vibrating microtome (Leica VT1000). Free-floating sections were blocked at 37°C for 1 hour in 0.3% Triton X-100 in PBS, containing 5% normal goat serum or fetal bovine serum. The slices were incubated with primary antibody overnight in 0.3% TritonX-100 in PBS, containing 5% normal goat serum or fetal bovine serum. Following washes, slices were incubated with corresponding fluorophore-conjugated secondaries for 1 hour at room temperature, washed and mounted (Fluoromount, Invitrogen). Antibodies and concentrations are given in **Table 2**.

**Table 2:** Primary and secondary antibodies for immunofluorescence.

| Specificity | Clone/Species | Conjugate | Vendor | Catalog Number | Purpose | Dilution |
| --- | --- | --- | --- | --- | --- | --- |
| DAT | Polyclonal/Rat | N/A | EMD Millipore Corp. | MAB369 | IHC | 1:500 |
| TH | Polyclonal/Rabbit<br>Polyclonal/Chicken | N/A | Encor | RPCA-TH<br>CPCA-TH | IHC | 1:500 |
| CD3 | Polyclonal/Rat | N/A | BioLegend | 100201 | IHC | 1:1000 |
| ChAT | Polyclonal/Goat | N/A | EMD Millipore Corp. | AB144P | IHC | 1:200 |
| GFP | Monoclonal/Mouse | N/A | Invitrogen | A11122 | IHC | 1:500 |
| mCherry | Monoclonal/Mouse | N/A | Encor | MPCA-mCherry | IHC | 1:500 |
| cFos | Polyclonal/Rabbit | N/A | Encor | RPCA-c-FOS | IHC | 1:500 |
| B220 | Polyclonal/Rat | N/A | BioLegend | 103202 | IHC | 1:500 |
| Rat | Polyclonal/Goat | Alexafluor-488<br>Alexafluor-647 | Invitrogen | A11006<br>A21247 | IHC | 1:500 |
| Rabbit | Polyclonal/Goat | Alexafluor-488<br>Alexafluor-568 | Invitrogen | A11034<br>A11011 | IHC | 1:500 |
| Guinea Pig | Polyclonal/Goat | Alexafluor-488 | Invitrogen | A11073 | IHC | 1:500 |
| Mouse | Polyclonal/Goat | Alexafluor-647 | Invitrogen | A21236 | IHC | 1:500 |
| Chicken | IgY | Alexafluor-647 | Invitrogen | A21449 | IHC | 1:1000 |
| Goat | Polyclonal/Donkey | Alexafluor-647 | Invitrogen | A21447 | IHC | 1:500 |

### Flow cytometry

Following terminal isoflurane anesthesia, the spleen was removed and placed in PBS on ice. Sterile tissue forceps were used to remove the spleen, place it on a slide and disrupt it to produce a splenocyte suspension. Slides were washed using 3 volumes of PBS to collect splenocyte suspension, followed by filtration through a 40 μm cell strainer placed over a 50mL conical tube. Following centrifugation (300 x g, 5 min), the cell pellet was resuspended in 70% Percoll density medium, and then underlaid below 37% Percoll. Centrifugation for 30 min at 500 x g with brakes and acceleration set to low results in a suspension of splenocytes at the interface between 37% and 70% Percoll. Cells were collected, washed twice with excess PBS, and then counted prior to the density being adjusted to 10,000 cells per microliter. Flow cytometry staining was conducted using 1 million cells suspended in 100uL of PBS. Antibodies and concentrations are given in **Table 3**.

**Table 3:** Antibodies for flow cytometry.

| Specificity | Clone/Species | Conjugate | Vendor | Catalog # | Purpose | Dilution |
| --- | --- | --- | --- | --- | --- | --- |
| CD3 | 17A2/Rat | Pac Blue | BioLegend | 100214 | FC | 1:50 |
| CD4 | GK1.5/Rat | AF700 | BioLegend | 100430 | FC | 1:100 |
| CD8a | QA17A07/Mouse | Spark 538 | BioLegend | 155020 | FC | 1:50 |
| CD127 | S18006K/Rat | PE/Fire 640 | BioLegend | 158213 | FC | 1:100 |
| KLRG1 | 2F1/KLRG1/Syrian Hamster | PE/Fire 810 | BioLegend | 138437 | FC | 1:200 |
| CD19 | 6D5/Rat | BV605 | BioLegend | 115540 | FC | 1:50 |
| CD45 | 30-F11/Rat | Spark Blue 574 | BioLegend | 103184 | FC | 1:200 |
| NK1.1 | S17016D/Mouse | Fire 700 | BioLegend | 156527 | FC | 1:100 |
| CD335 | 29A1.4/Rat | PerCP Fire 806 | BioLegend | 137650 | FC | 1:50 |
| CD11b | M1-70/Rat | FITC | BioLegend | 101206 | FC | 1:200 |
| F4/80 | BM8/Rat | Spark NIR 685 | BioLegend | 123168 | FC | 1:50 |
| CD44 | IM7/Rat | Spark Yg 594 | BioLegend | 103078 | FC | 1:100 |
| CD62L | MEL-14/Rat | BV711 | BioLegend | 104445 | FC | 1:200 |
| IgM | RMM-1/Rat | PE | BioLegend | 406507 | FC | 1:200 |
| IgD | 11-26c.2a/Rat | PerCP Cy5.5 | BioLegend | 405709 | FC | 1:200 |

## Acknowledgements

We acknowledge assistance from Kaitlyn N. Kiel and technical assistance from the University of Florida Center for Immunology and Transplantation.

---

**Supplementary Figure 1.**
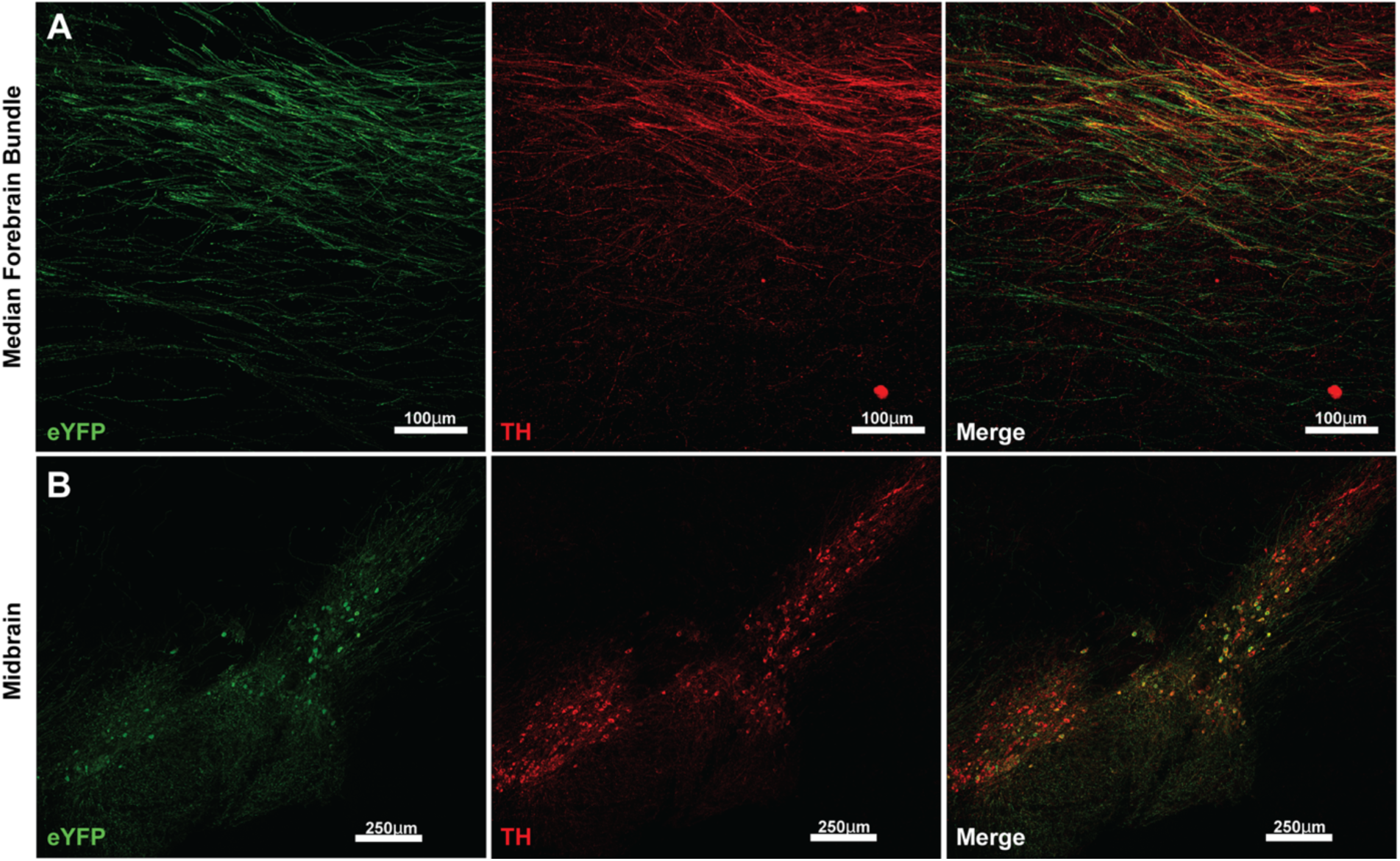
Single channel control images for EYFP-expressing midbrain and median forebrain bundle verify injection site and expression in midbrain dopamine neuron projections. **Panel A:** Median forebrain bundle of DAT-Cre mice injected with AAV2-Ef1a-DIO-EYFP shows EYFP+ fibers and overlap with TH immunoreactive fibers. **Panel B:** At the midbrain site of injection, EYFP+ cell bodies overlap with TH+ midbrain dopamine neurons.

**Supplementary Figure 2.**
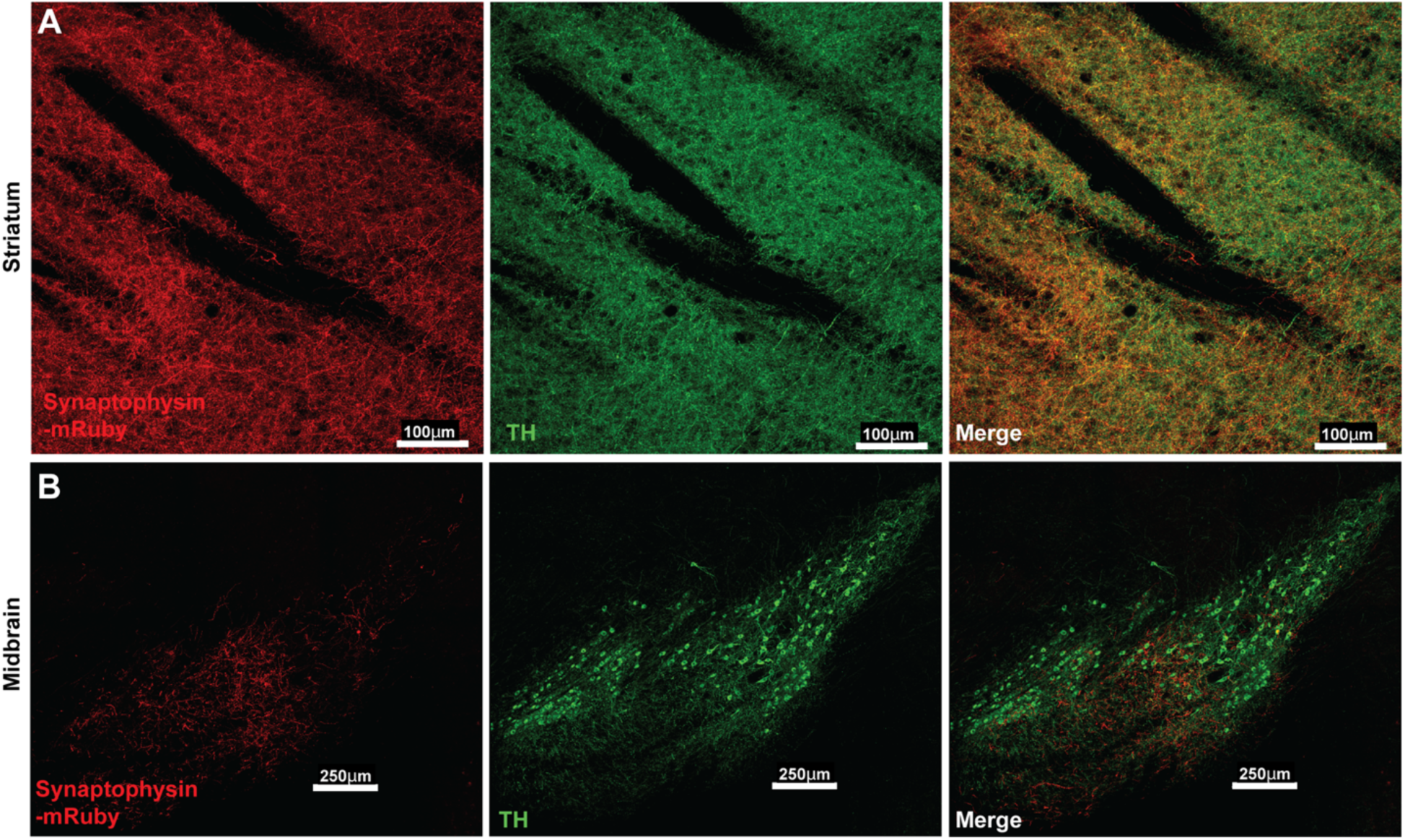
Single channel control images for synaptophysin-expressing midbrain and striatum verify injection site and expression in midbrain dopamine neuron projections. **Panel A:** Striatum of DAT-Cre mice injected with AAV1-hSyn-Flex mGFP-2A-Synaptophysin-mRuby shows synaptophysin-mRuby+ terminals and overlap with TH immunoreactive terminals. **Panel B**: At the midbrain site of injection, synaptophysin-mRuby+ cell bodies overlap with TH+ midbrain dopamine neurons.

**Supplementary Figure 3.**
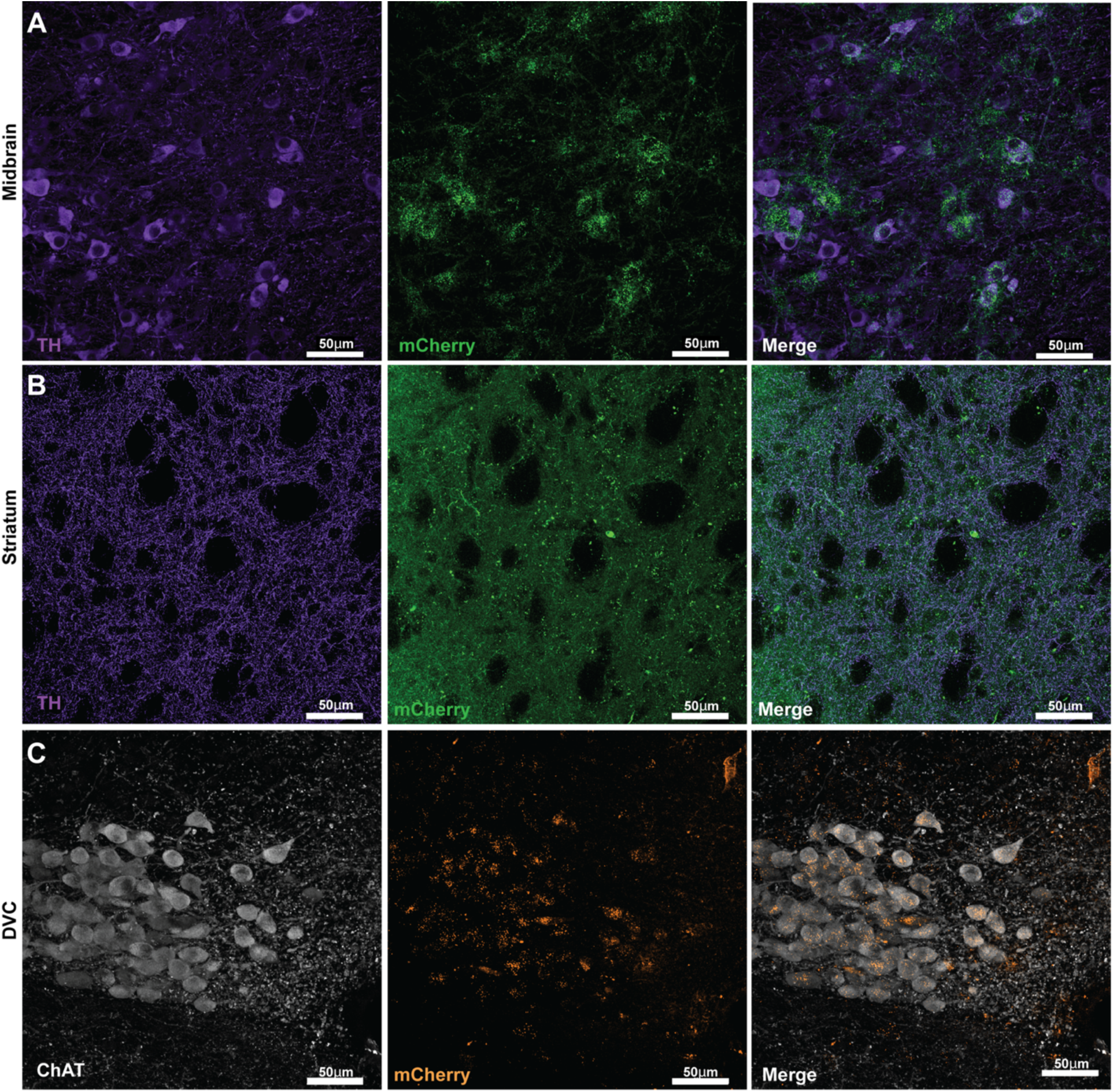
mCherry-tagged Cre-dependent Gq-DREADDs are expressed in midbrain dopamine neurons and terminal regions including the DVC. DAT-Cre mice received midbrain microinjections of Cre-dependent AAV-DIO-mCherry-hM3D (Gq-DREADD), or AAV2-hSyn-DIO-mCherry (empty vector). mCherry fluorescence was assessed in midbrain and in terminal regions. **Panel A**: Single channel control images show midbrain dopamine neurons, identified by expression of TH, express mCherry-tagged Gq-DREADDs. **Panel B**: mCherry^+^ terminals are evident throughout the striatum, indicating this AAV construct expresses Gq-DREADD receptors in terminal regions. **Panel C**: mCherry^+^ Gq-DREADD-expressing terminals are evident throughout the DVC.

**Supplementary Figure 4:**
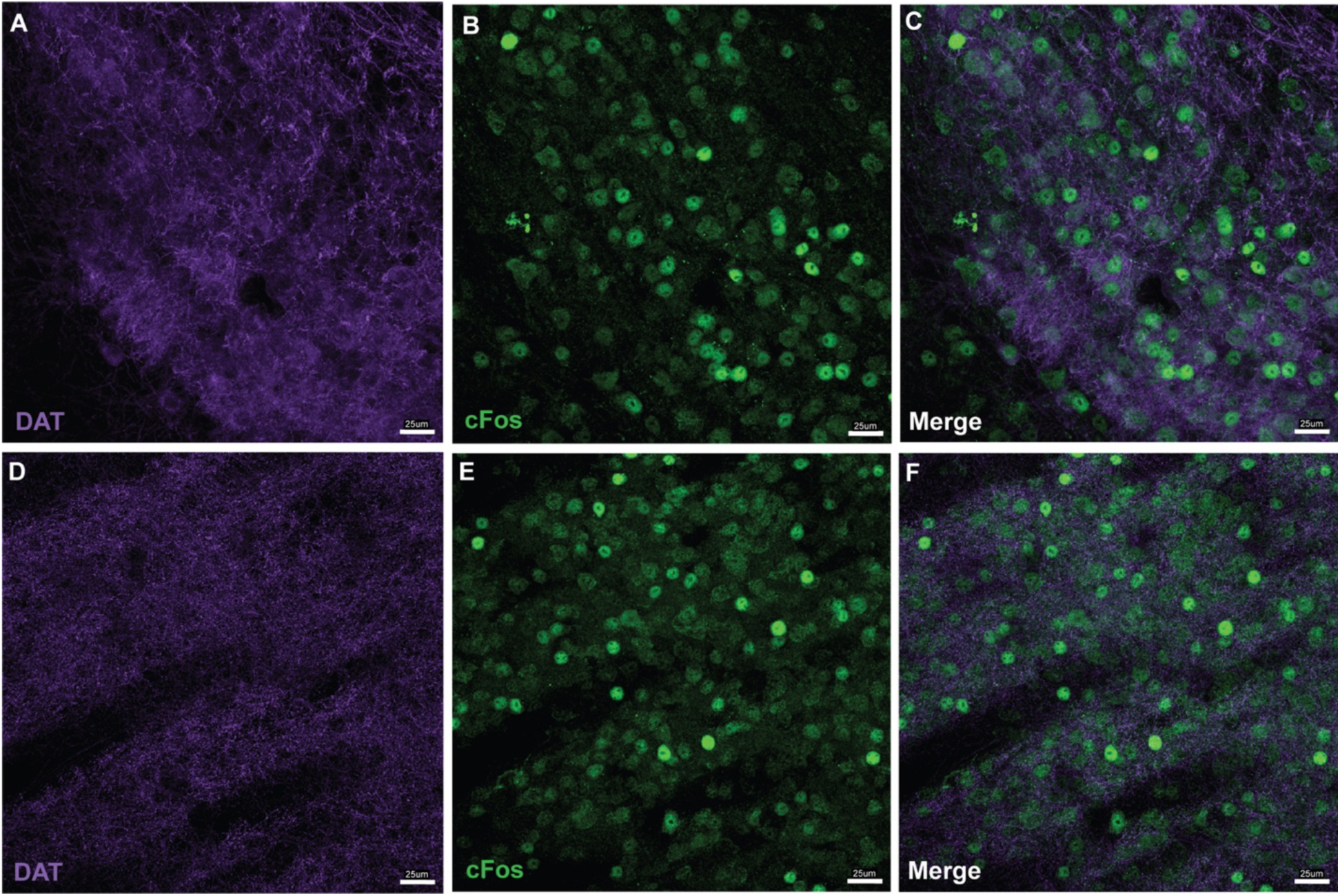
Gq-DREADD stimulation of midbrain dopamine neurons increases cFos expression in both midbrain and striatum. DAT-Cre mice received midbrain microinjections of AAV-hSyn-DIO-mCherry-hM3D (Gq-DREADD), or AAV2-hSyn-DIO-mCherry (empty vector). **Panel A-C:** Midbrain dopamine neurons of mice that received excitatory Gq-DREADDs, identified by expression of DAT, show robust cFos expression following systemic CNO administration. **Panel D-F:** Striatal terminal regions of dopamine neurons, identified by DAT expression, also exhibit robust cFos expression.

**Supplementary Figure 5:**
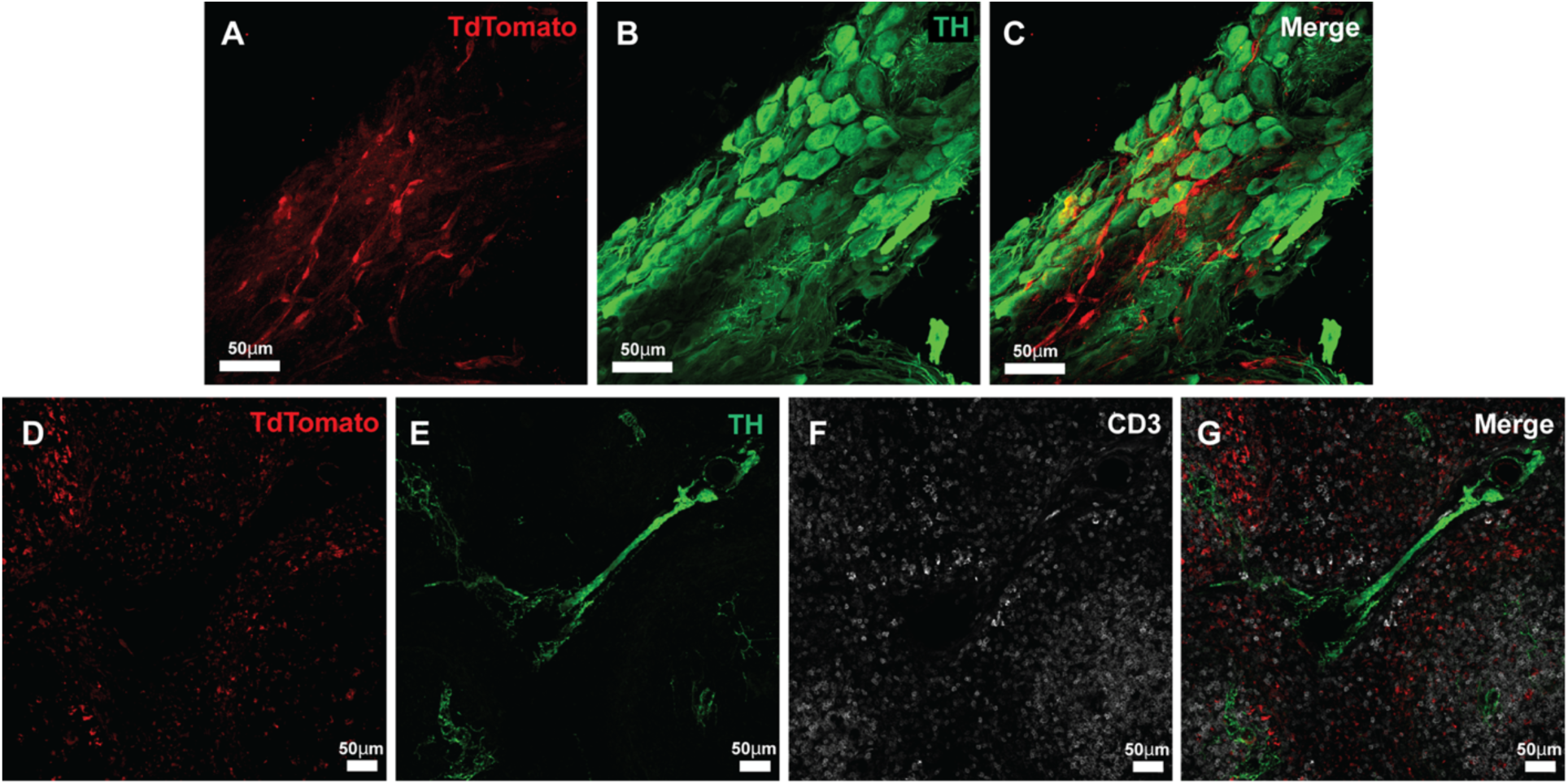
Single channel control images for TdTomato+ signals in the celiac ganglion and spleen. **Panel A-C**: Single channel images showing TdTomato (Panel A), TH (Panel B) and a merge image (Panel C) showing TdTomato+ cells in proximity to TH+ cell bodies in the celiac ganglion of Ai9 mice that received midbrain injections of pENN-AAV1-hSyn-Cre-WPRE-hGH. **Panel D-G**: Single channel images showing TdTomato expression (Panel D), TH-expressing fibers (Panel E) and CD3+ T-cell zones (Panel F), and a merged image (Panel G) showing TdTomato+ fibers surrounding T-cell zones in the spleen of Ai9 mice that received midbrain injections of pENN-AAV1-hSyn-Cre-WPRE-hGH.

**Supplementary Figure 6:**
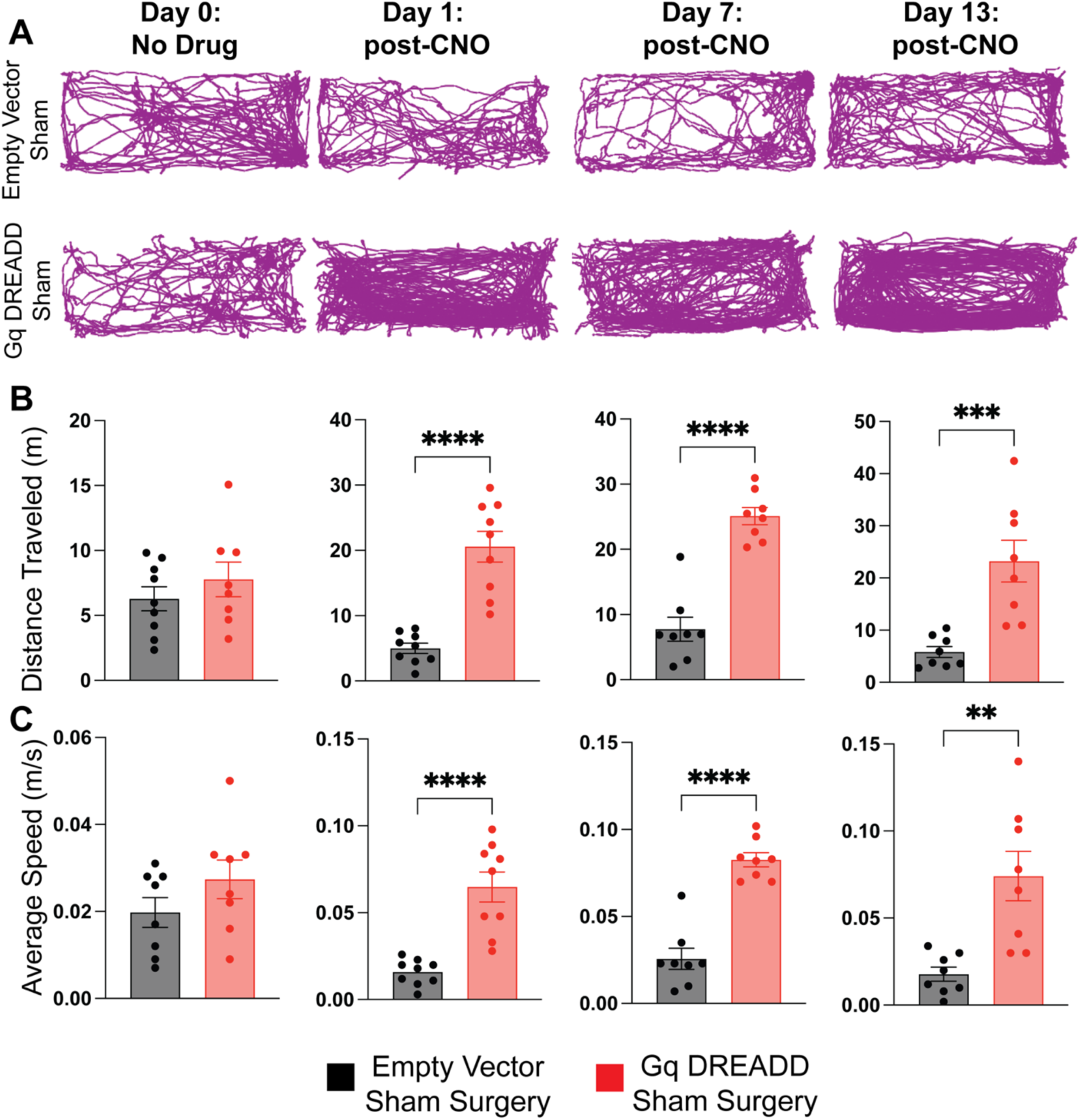
Gq-DREADD stimulation results in increased locomotor activity following intraperitoneal injection of 1mg/kg clozapine-*N*-oxide, in animals without vagotomy. **Panel A:** Locomotor activity in the home cage was measured prior to drug injection (Day 0), and on Days 1, 7 and 13 during the CNO administration period. **Panel B:** On Day 0, total distance traveled is not significantly different between animals receiving Gq-DREADD in midbrain dopamine neurons relative to empty vector (AAV2-hSyn-DIO-mCherry). Consistent with the literature in the field, CNO increases locomotor activity in animals with midbrain dopamine neuron Gq-DREADDs compared to empty vector on Days 1, 7 and 13. **Panel C:** Average speed is also significantly increased in animals receiving Gq-DREADDs relative to empty vector control (AAV2-hSyn-DIO-mCherry) on Days 1, 7 and 13. *(n=8 independent biological replicates; ***p<0.005,****p<0.0005, two-tailed unpaired Student’s T-test).* See also **Supplementary Movie 1**.

**Supplementary Figure 7:**
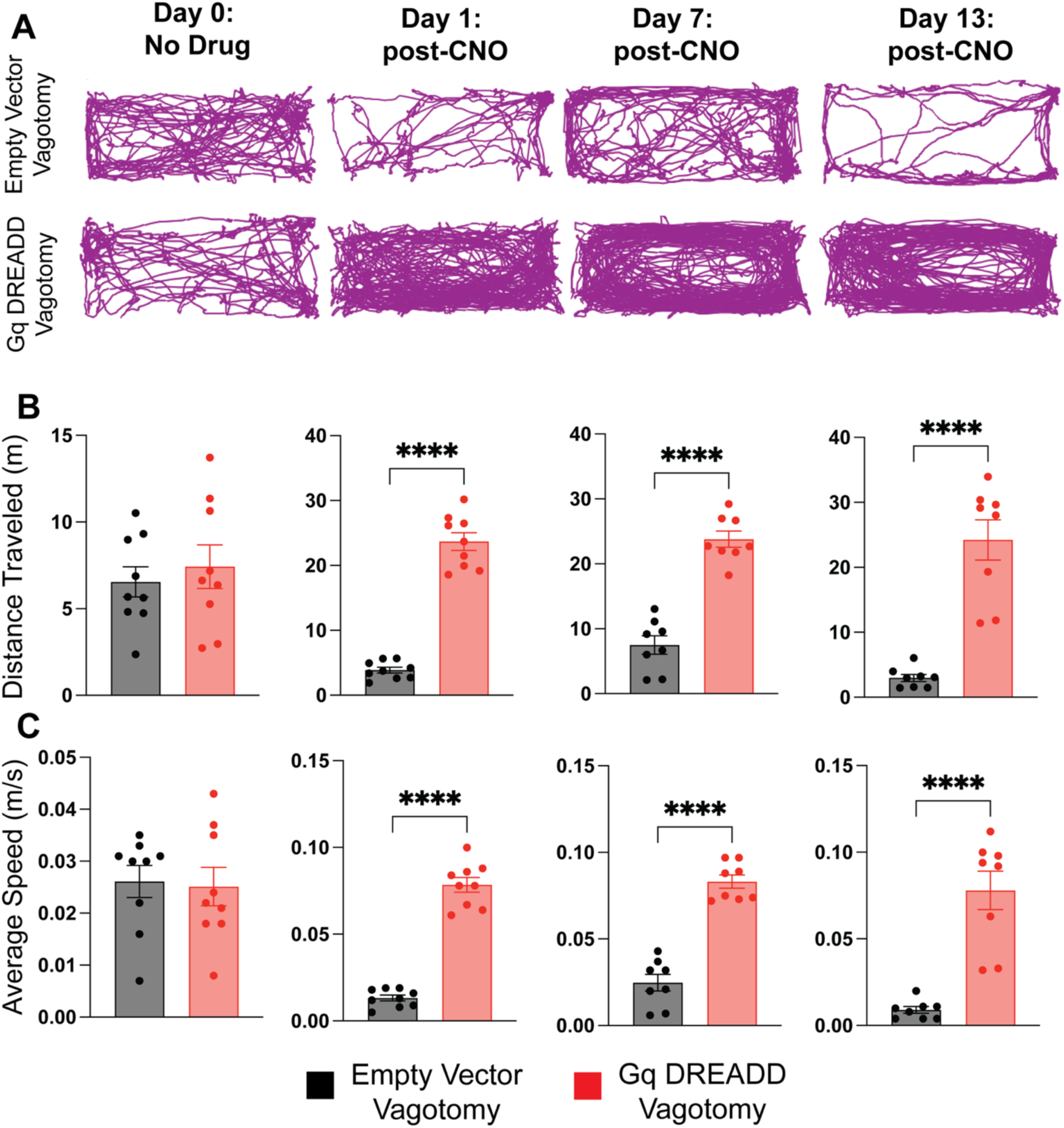
Following cervical vagotomy, Gq-DREADD stimulation results in increased locomotor activity following intraperitoneal injection of 1mg/kg clozapine-*N*-oxide. **Panel A:** Locomotor activity in the home cage was measured prior to drug injection (Day 0), and on Days 1, 7 and 13 during the CNO administration period. **Panel B:** On Day 0, total distance traveled is not significantly different between animals receiving Gq-DREADD in midbrain dopamine neurons relative to empty vector (AAV2-hSyn-DIO-mCherry). Consistent with the literature in the field, CNO increases locomotor activity in animals with midbrain dopamine neuron Gq-DREADDs compared to empty vector on Days 1, 7 and 13. **Panel C:** Average speed is also significantly increased in animals receiving Gq-DREADDs relative to empty vector control (AAV2-hSyn-DIO-mCherry) on Days 1, 7 and 13. *(n=8 independent biological replicates; ***p<0.005,****p<0.0005, two-tailed unpaired Student’s T-test).* See also **Supplementary Movie 2**.

**Supplementary Figure 8:**
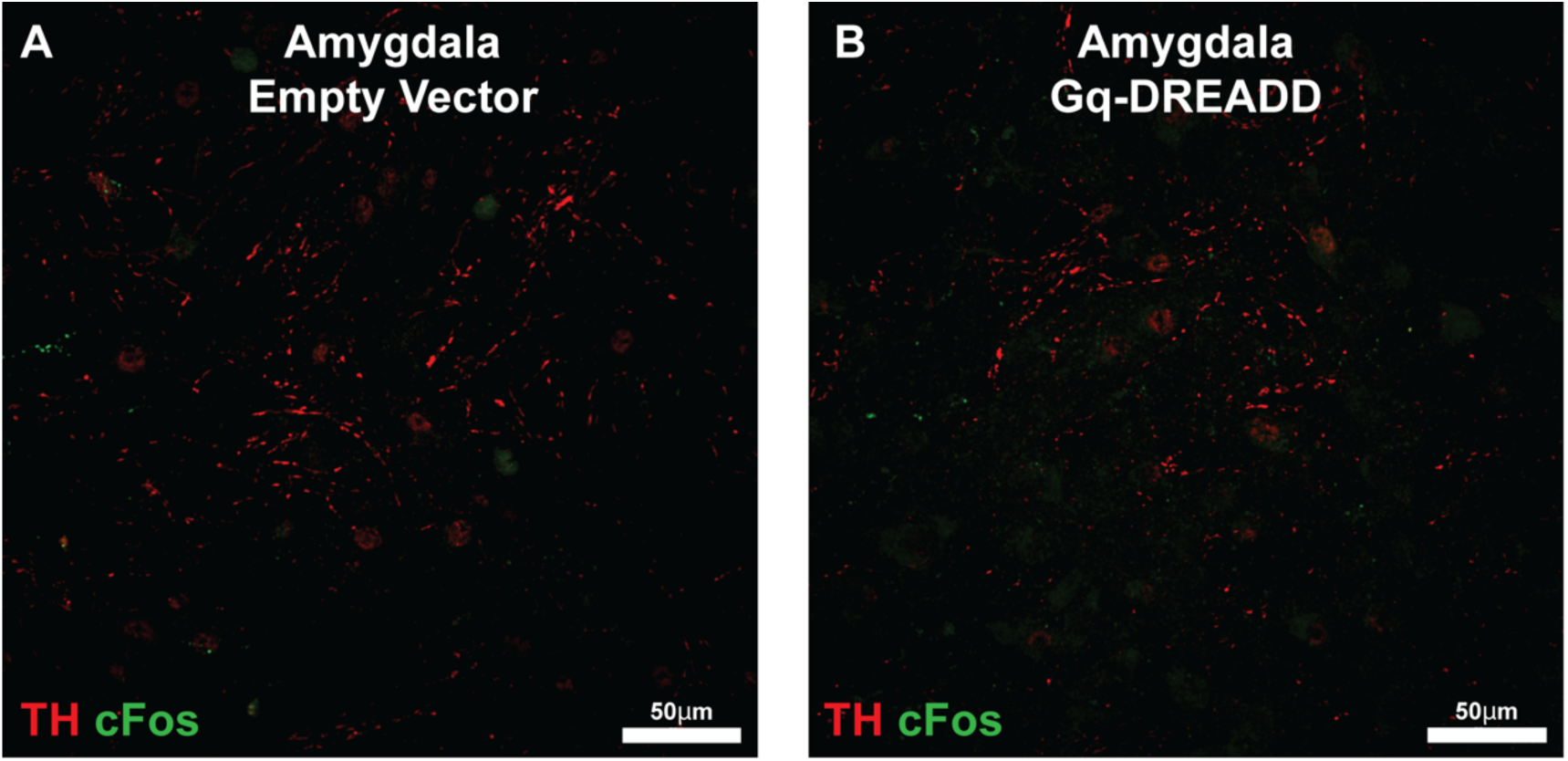
Chemogenetic activation of midbrain dopamine neurons does not increase cFos expression in amygdala. **Panels A-B:** In the central amygdala, identified by anatomical landmarks and presence of TH+ fibers, chemogenetic activation of midbrain dopamine neurons does not increase cFos expression.

**Supplementary Figure 9.**
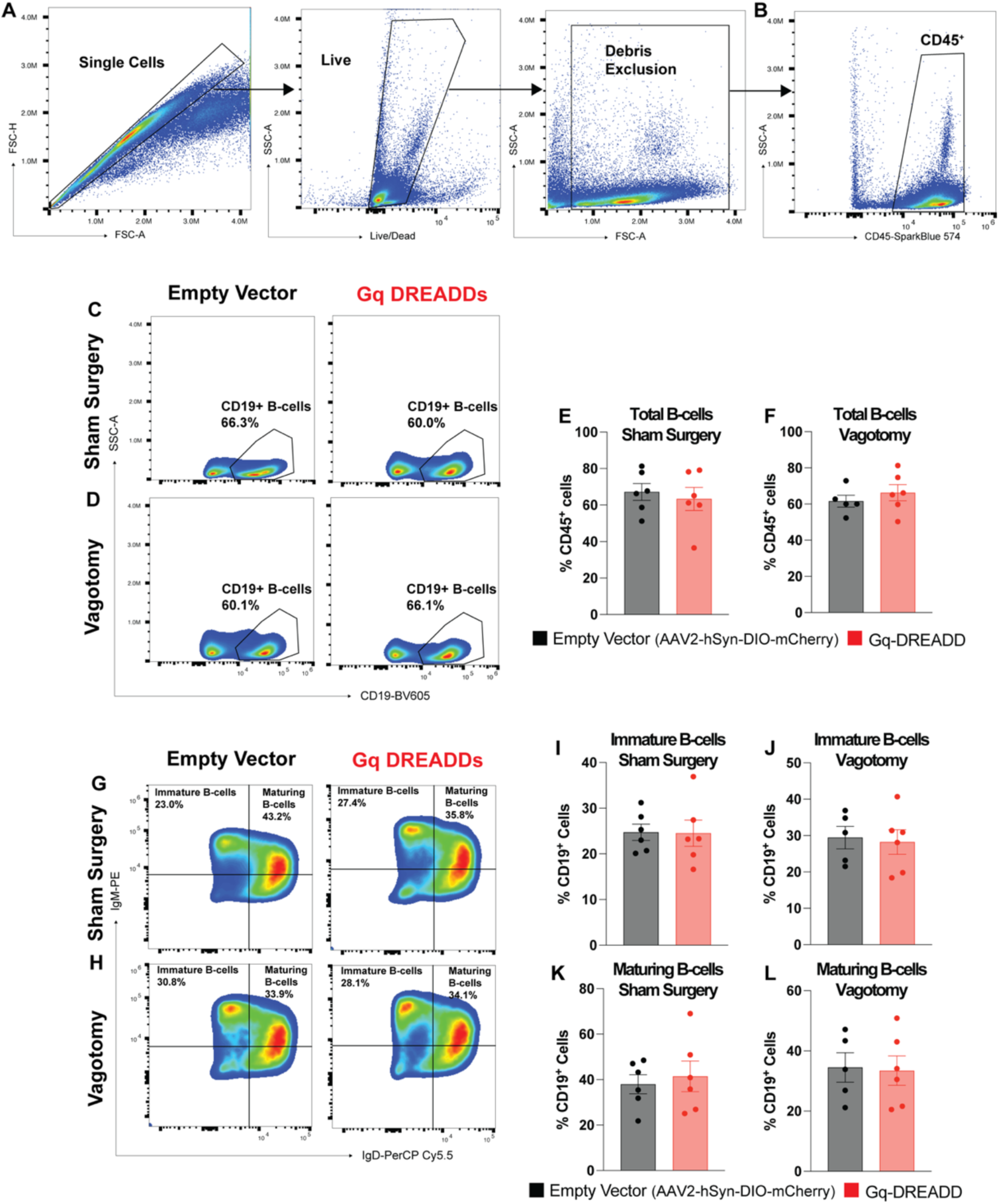
B-cell populations are not significantly altered when midbrain dopamine neurons are stimulated with Gq DREADDs. **Panel A-B:** Single, live cells, free of debris were gated for expression of CD45. **Panel C-D**: Total CD19+ B-cells, and **Panels G-H**: Immature and Maturing B-cells, identified by expression of IgM alone or co-expression of IgM and IgD, were assessed to determine impact of midbrain dopamine neuron activation. **Panel E-F**: Total CD19+ B-cells were not significantly altered when midbrain dopamine neurons were activated. **Panel I-L**: Immature B-cells and Maturing B-cells were not significantly altered when midbrain dopamine neurons were activated. *(n=6 independent biological replicates; two-tailed unpaired Student’s T-test, p>0.05; N.S.)*

**Supplementary Figure 10.**
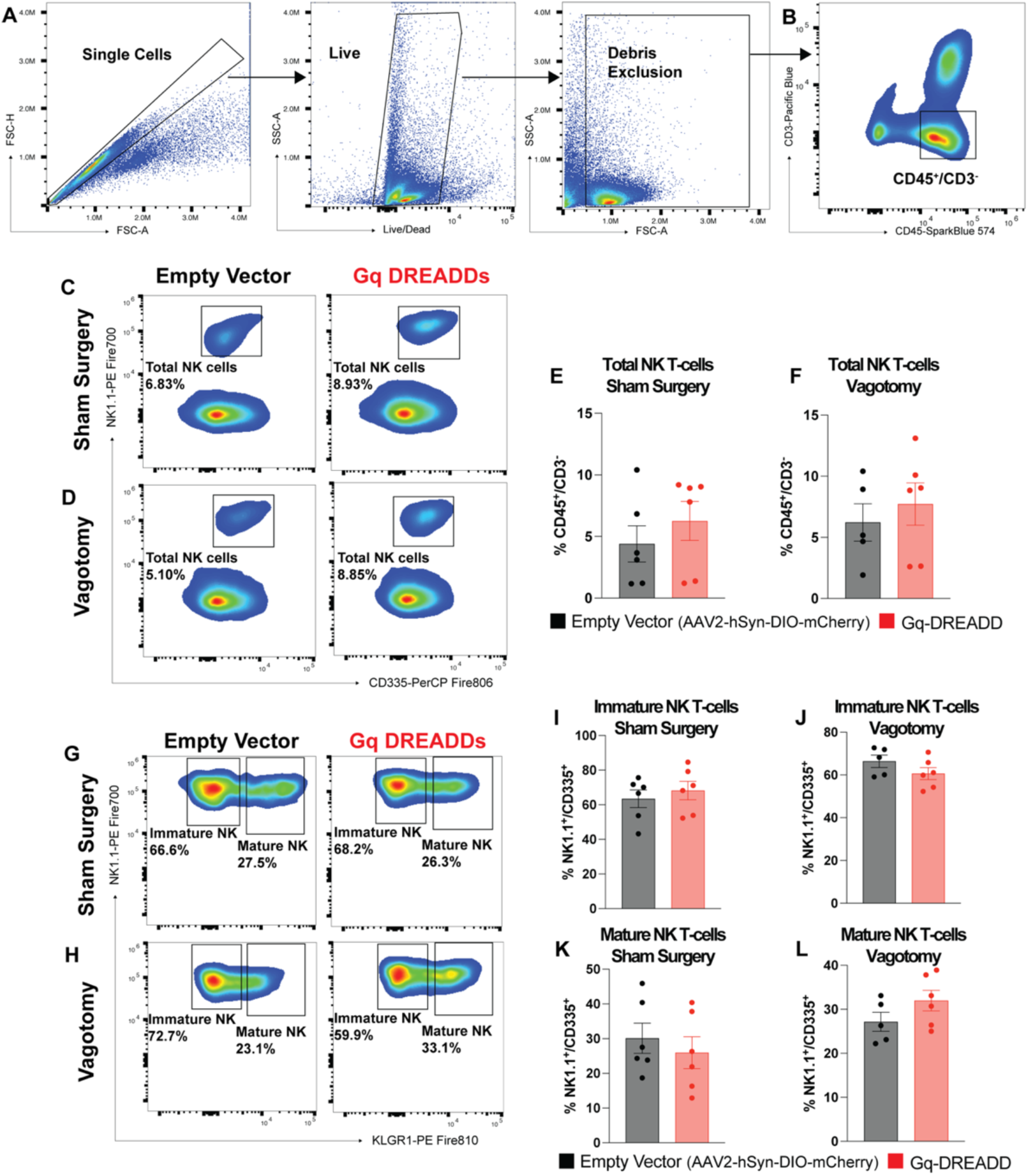
NK-cell populations are not significantly altered when midbrain dopamine neurons are stimulated with Gq DREADDs. **Panel A-B:** Single, live cells, free of debris were gated for expression of CD45 and lack of CD3 expression (CD45+/CD3-) to identify NK cells. **Panel C-D**: Total NK cells, identified by expression of CD335 and NK1.1, and **Panel G-H**: Immature and mature NK cells, identified by expression of KLGR1, were assessed to determine impact of midbrain dopamine neuron activation. **Panel E-F**: Total NK-cells were not significantly altered when midbrain dopamine neurons were activated. **Panel I-L**: Immature and Mature NK-cells were not significantly altered when midbrain dopamine neurons were activated. *(n=6 independent biological replicates; two-tailed unpaired Student’s T-test, p>0.05; N.S.)*

**Supplementary Figure 11.**
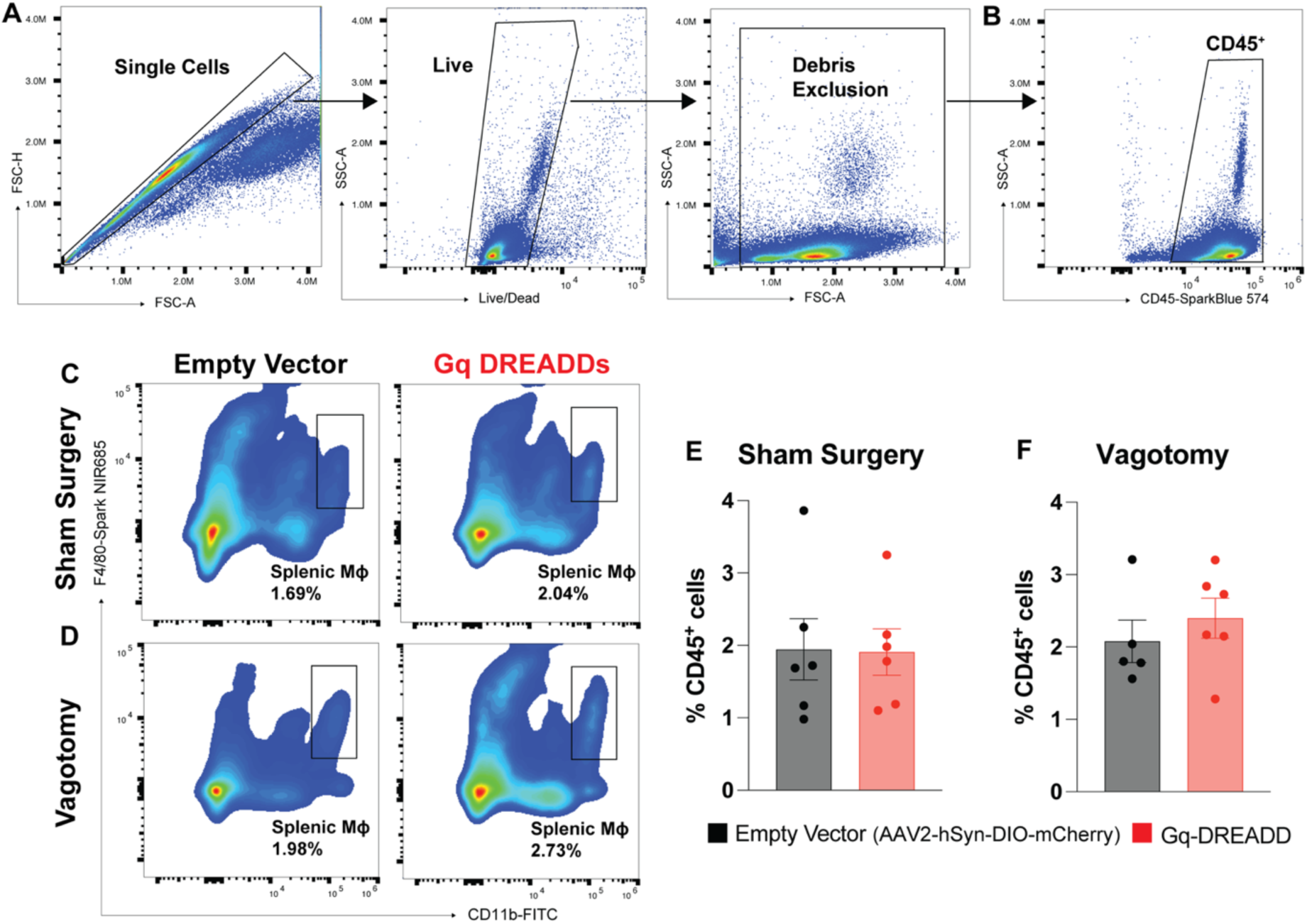
Splenic macrophages are not significantly altered when midbrain dopamine neurons are stimulated with Gq DREADDs. **Panel A-B:** Single, live cells, free of debris were gated for CD45 expression. **Panel C-D**: Splenic macrophages were identified by co-expression of CD11b and F4/80 (CD11b+/F480+). Panel E-F: Splenic macrophages were not significantly altered when midbrain dopamine neurons were activated. *(n=6 independent biological replicates; two-tailed unpaired Student’s T-test, p>0.05; N.S.)*.

**Supplementary Figure 12.**
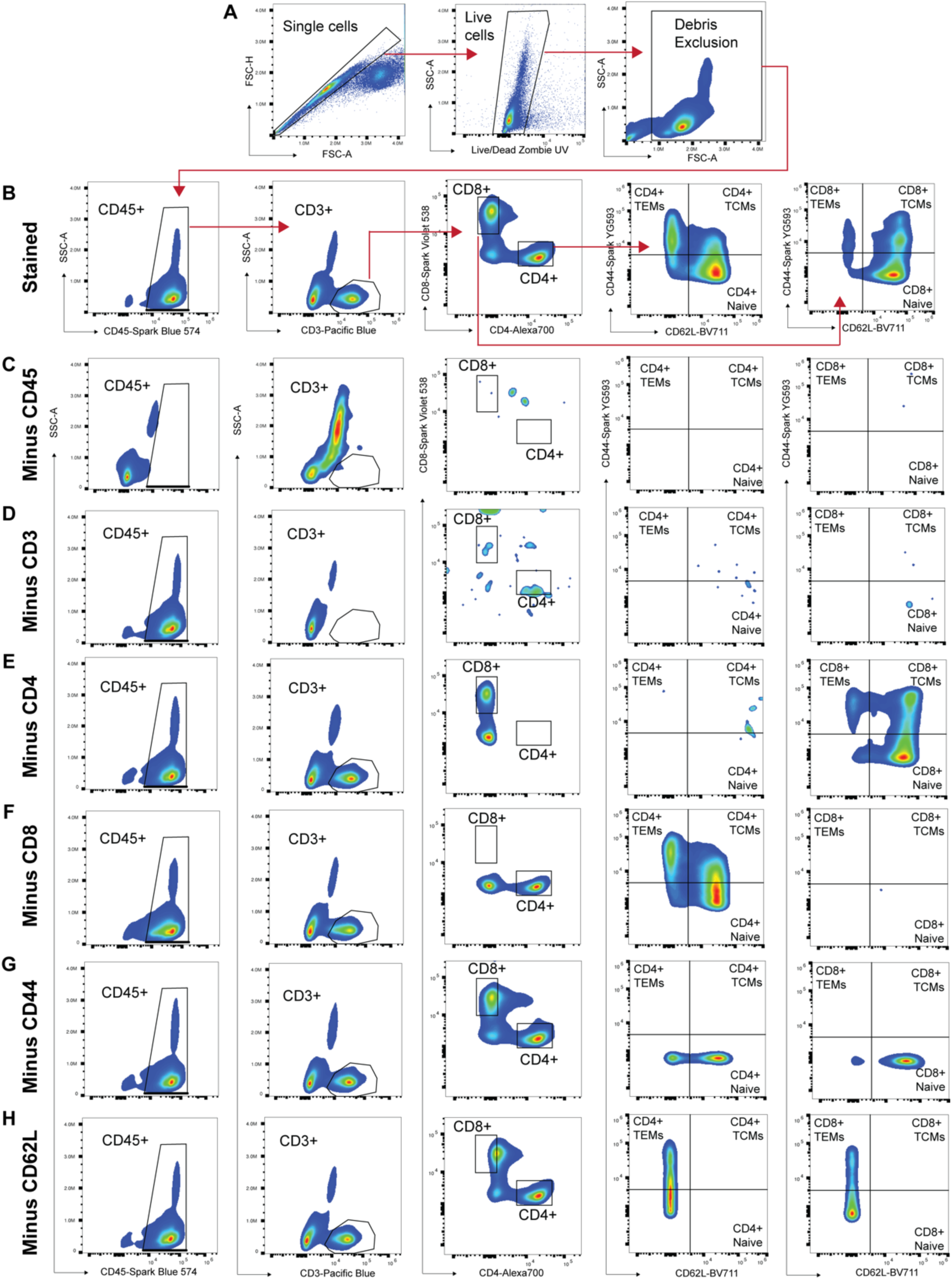
Fluorescence-minus-one generated gating strategy to analyze splenic T-cell populations. **Panel A**: Single, live cells free of debris were (**Panel B**) assessed for expression of CD45. **Panel B**: Subsequently, cells expressing CD3 were assessed for CD4 and CD8 expression and then assessed for T-central memory (TCM), T-effector memory (TEM) and Naïve T-cells. **Panels C-H**: Gates for analyses for each of the above populations were set by fluorescence-minus-one gating strategy.

**Supplementary Figure 13.**
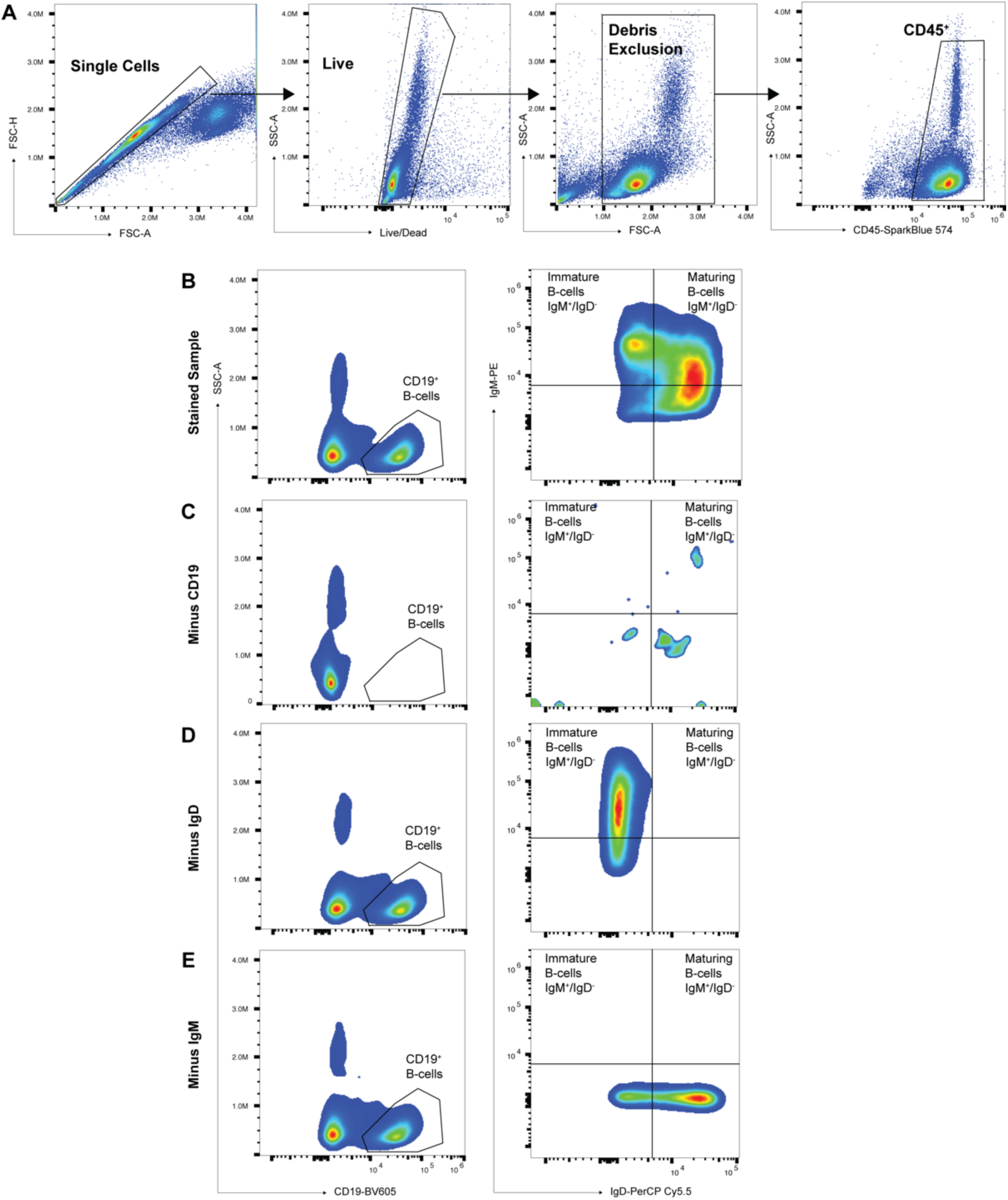
Fluorescence-minus-one generated gating strategy to analyze splenic B-cell populations. **Panel A**: Single, live cells free of debris were (**Panel B**) assessed for expression of CD45. **Panel B**: Subsequently, cells expressing CD19 were assessed for IgM and IgD expression. **Panels C-E**: Gates for analyses for each of the above populations were set by fluorescence-minus-one gating strategy.

**Supplementary Figure 14.**
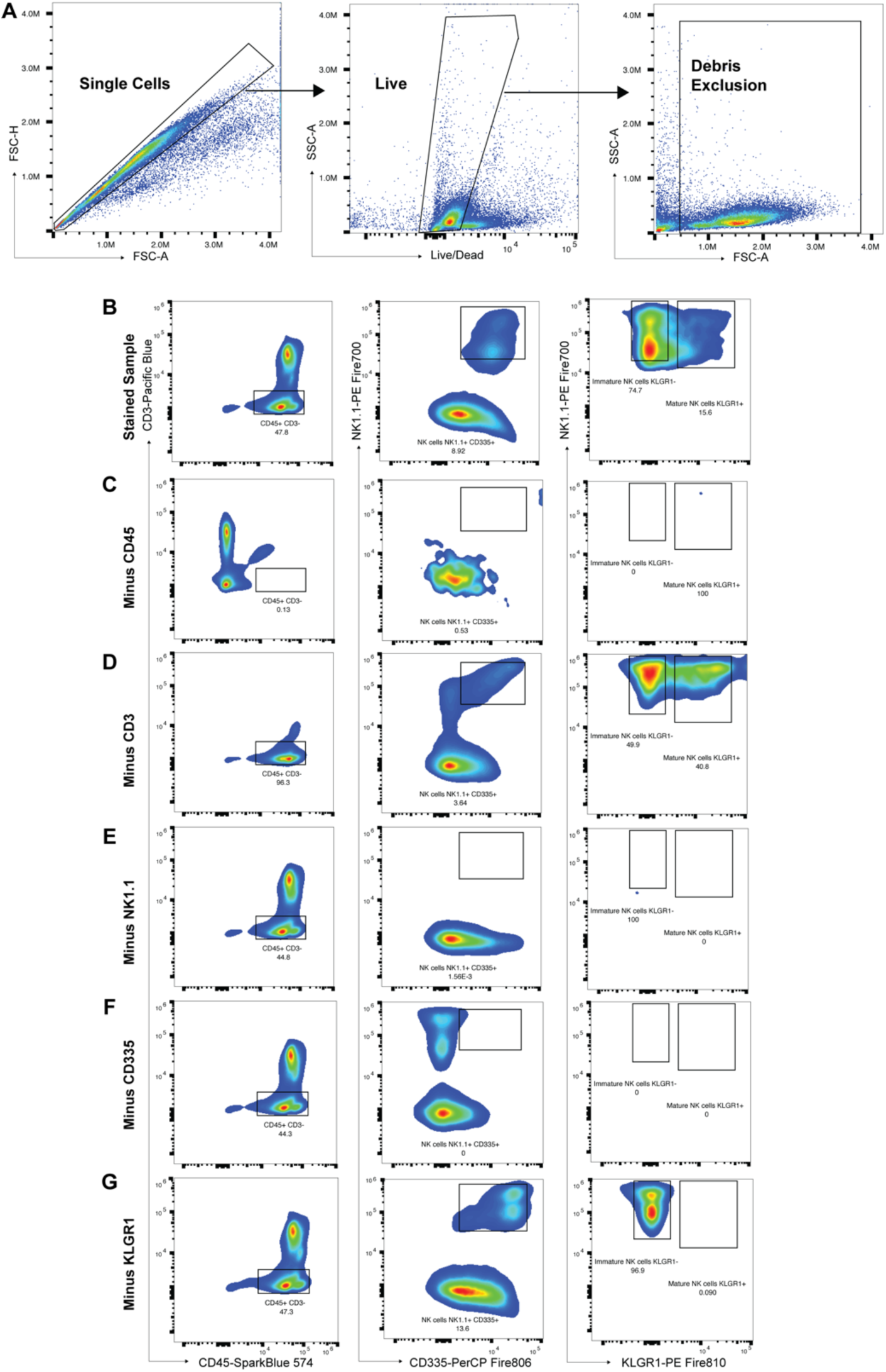
Fluorescence-minus-one generated gating strategy to analyze splenic NK-cell populations. **Panel A**: Single, live cells free of debris were (**Panel B**) assessed for expression of CD45 without CD3 expression (CD45+/CD3-). **Panel B**: Subsequently, cells co-expressing CD335 and NK1.1 (total NK-cells) were assessed for KLGR1 expression. **Panels C-G**: Gates for analyses for each of the above populations were set by fluorescence-minus-one gating strategy.

**Supplementary Figure 15.**
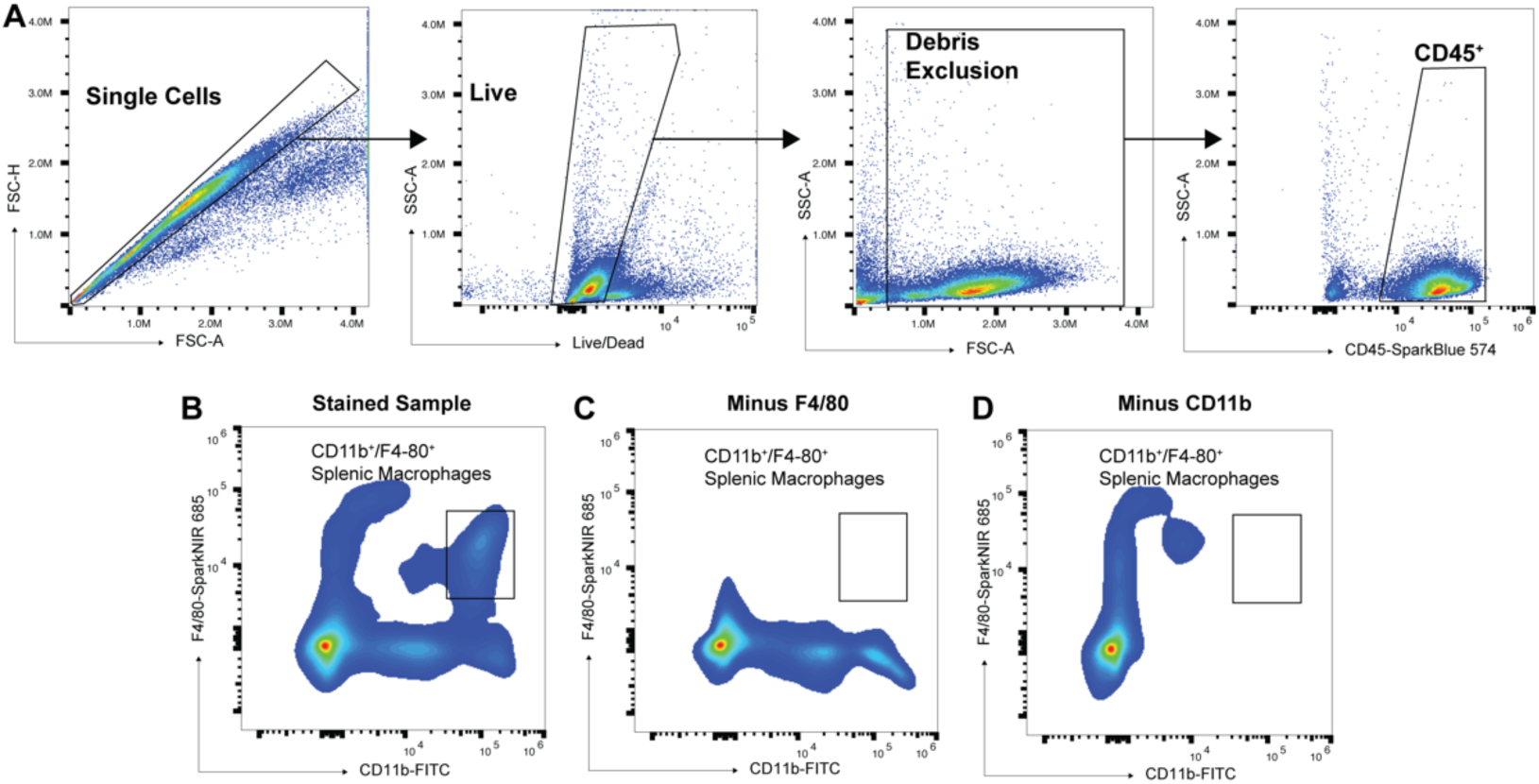
Fluorescence-minus-one generated gating strategy to analyze splenic macrophages. **Panel A**: Single, live cells free of debris were (**Panel B**) assessed for expression of CD45. **Panel B**: Subsequently, cells were assessed for co-expression of CD11b and F4/80 (splenic macrophages). **Panels C-D**: Gates for analyses for each of the above populations were set by fluorescence-minus-one gating strategy.

**Supplementary Video 1: Increased locomotor activity evident throughout 14 days of systemic CNO treatment in mice that received excitatory Gq-DREADD AAV injection in midbrain dopamine neurons and sham cervical surgery.**

**Supplementary Video 2: Increased locomotor activity evident throughout 14 days of systemic CNO treatment in mice that received excitatory Gq-DREADD AAV injection in midbrain dopamine neurons and cervical vagotomy.**

