## Supplementary Table 1 for "Midbrain dopamine neuronal activity modulates splenic immunity through a brain-to-body circuit"

### Supplementary Tables 1

Intrinsic properties (baseline) – Supplementary table to Figure 3B

| | GFP+ | | GFP- | | p value |
|---|---|---|---|---|---|
| | Average | ±SEM | Average | ±SEM | |
| Ri (MΩ) | 639.64 | 100.51 | 878.89 | 80.26 | 0.0764 |
| τ (msec) | 0.46 | 0.03 | 0.36 | 0.03 | 0.6094 |
| Vthresh (mV) | -32.11 | 1.50 | -31.90 | 1.64 | 0.9256 |
| Ithresh (pA) | 81.82 | 8.42 | 50.00 | 0.00 | **0.0012** |
| V height (mV) | 70.61 | 2.29 | 59.03 | 2.19 | **0.0008** |
| APhalfwidth (msec) | 0.73 | 0.04 | 0.55 | 0.02 | **0.0003** |
| f/I slope (Hz/pA) | 0.08 | 0.01 | 0.25 | 0.05 | **0.0013** |
| SFA ratio (mean $2^{nd}/3^{rd}$) | 1.07 | 0.11 | 0.59 | 0.05 | **0.0006** |

Passive properties in response to dopamine – Supplementary table to Figure 3D

| | GFP+ | | | | | GFP- | | | | |
|---|---|---|---|---|---|---|---|---|---|---|
| | Before | | After | | p value | Before | | After | | p value |
| | Average | ±SEM | Average | ±SEM | | Average | ±SEM | Average | ±SEM | |
| Ri (MΩ) | 639.64 | 100.51 | 431.09 | 47.79 | 0.07 | 878.89 | 80.26 | 656.37 | 63.43 | **0.036** |
| τ (msec) | 0.46 | 0.03 | 0.50 | 0.03 | 0.43 | 0.43 | 0.07 | 0.43 | 0.03 | 0.946 |

Active properties in response to dopamine – Supplementary table to Figure 3G-I

| | GFP+ | | | | |
|---|---|---|---|---|---|
| | Before | | After | | p value |
| | Average | ±SEM | Average | ±SEM | |
| Vthresh (mV) | -32.11 | 1.50 | 27.84 | 1.12 | **0.028** |
| Ithresh (pA) | 81.82 | 8.42 | 143.18 | 15.53 | **0.001** |
| V height (mV) | 70.61 | 2.29 | 63.38 | 2.29 | **0.031** |
| APhalfwidth (msec) | 0.73 | 0.04 | 0.72 | 0.03 | 0.852 |
| f/I slope (Hz/pA) | 0.08 | 0.01 | 0.07 | 0.01 | 0.516 |
| SFA ratio (mean $2^{nd}/3^{rd}$) | 1.07 | 0.11 | 1.01 | 0.08 | 0.706 |

| | Responsive GFP- | | | | |
|---|---|---|---|---|---|
| | Before | | After | | p value |
| | Average | ±SEM | Average | ±SEM | |
| Vthresh (mV) | -30.56 | 2.53 | -29.31 | 2.69 | 0.740 |
| Ithresh (pA) | 50.00 | 0.00 | 85.00 | 13.02 | **0.015** |
| V height (mV) | 61.14 | 2.85 | 55.04 | 3.52 | 0.195 |
| APhalfwidth (msec) | 0.56 | 0.02 | 0.56 | 0.02 | 0.940 |
| f/I slope (Hz/pA) | 0.15 | 0.03 | 0.08 | 0.02 | 0.051 |
| SFA ratio (mean $2^{nd}/3^{rd}$) | 0.53 | 0.07 | 0.54 | 0.07 | 0.941 |

| | Unresponsive GFP- | | | | |
|---|---|---|---|---|---|
| | Before | | After | | p value |
| | Average | ±SEM | Average | ±SEM | |
| Vthresh (mV) | -33.39 | 2.04 | -36.30 | 2.57 | 0.388 |
| Ithresh (pA) | 50.00 | 0.00 | 50.00 | 0.00 | NA |
| V height (mV) | 56.69 | 3.37 | 55.38 | 2.96 | 0.774 |
| APhalfwidth (msec) | 0.53 | 0.03 | 0.52 | 0.03 | 0.863 |
| f/I slope (Hz/pA) | 0.35 | 0.09 | 0.33 | 0.08 | 0.899 |
| SFA ratio (mean $2^{nd}/3^{rd}$) | 0.66 | 0.08 | 0.69 | 0.09 | 0.841 |

Alpha synapse – Supplementary to Figure 3J

| | GFP+ | | | | | | | | | |
|---|---|---|---|---|---|---|---|---|---|---|
| | Pulse 1 | | Pulse 2 | | Pulse 3 | | Pulse 4 | | Pulse 5 | |
| | Average | ±SEM | Average | ±SEM | Average | ±SEM | Average | ±SEM | Average | ±SEM |
| Before | 1.0 | 0.0 | 1.26 | 0.03 | 1.32 | 0.05 | 1.35 | 0.06 | 1.36 | 0.06 |
| After | 1.0 | 0.0 | 1.17 | 0.03 | 1.21 | 0.04 | 1.22 | 0.05 | 1.22 | 0.05 |
| p value | NA | | **0.048** | | 0.097 | | 0.071 | | 0.067 | |

| | Responsive GFP- | | | | | | | | | |
|---|---|---|---|---|---|---|---|---|---|---|
| | Pulse 1 | | Pulse 2 | | Pulse 3 | | Pulse 4 | | Pulse 5 | |
| | Average | ±SEM | Average | ±SEM | Average | ±SEM | Average | ±SEM | Average | ±SEM |
| Before | 1.0 | 0.0 | 1.26 | 0.02 | 1.34 | 0.03 | 1.37 | 0.04 | 1.37 | 0.05 |
| After | 1.0 | 0.0 | 1.16 | 0.03 | 1.19 | 0.03 | 1.19 | 0.04 | 1.18 | 0.04 |
| p value | NA | | **0.008** | | **0.004** | | **0.004** | | **0.005** | |

| | Unresponsive GFP- | | | | | | | | | |
|---|---|---|---|---|---|---|---|---|---|---|
| | Pulse 1 | | Pulse 2 | | Pulse 3 | | Pulse 4 | | Pulse 5 | |
| | Average | ±SEM | Average | ±SEM | Average | ±SEM | Average | ±SEM | Average | ±SEM |
| Before | 1.0 | 0.0 | 1.27 | 0.05 | 1.37 | 0.09 | 1.41 | 0.11 | 1.45 | 0.12 |
| After | 1.0 | 0.0 | 1.20 | 0.04 | 1.24 | 0.06 | 1.26 | 0.07 | 1.28 | 0.07 |
| p value | NA | | 0.338 | | 0.258 | | 0.259 | | 0.250 | |
